# FilaBuster: A Strategy for Rapid, Specific, and Spatiotemporally Controlled Intermediate Filament Disassembly

**DOI:** 10.1101/2025.04.20.649718

**Authors:** Andrew S. Moore, Tommy Krug, Simon B. Hansen, Alexander V. Ludlow, Jonathan B. Grimm, Anthony X. Ayala, Sarah E. Plutkis, Nan Wang, Robert D. Goldman, Ohad Medalia, Luke D. Lavis, David A. Weitz, Jennifer Lippincott-Schwartz

## Abstract

Intermediate filaments (IFs) play key roles in cellular mechanics, signaling, and organization, but tools for their rapid, selective disassembly remain limited. Here, we introduce FilaBuster, a photochemical approach for efficient and spatiotemporally controlled IF disassembly in living cells. FilaBuster uses a three-step strategy: (1) targeting HaloTag to IFs, (2) labeling with a covalent photosensitizer ligand, and (3) light-induced generation of localized reactive oxygen species to trigger filament disassembly. This modular strategy applies broadly across IF subtypes—including vimentin, GFAP, desmin, peripherin, and keratin 18—and is compatible with diverse dyes and imaging platforms. Using vimentin IFs as a model system, we establish a baseline implementation in which vimentin-HaloTag labeled with a photosensitizer HaloTag ligand triggers rapid and specific IF disassembly upon light activation. We then refine this approach by (i) expanding targeting strategies to include a vimentin nanobody-HaloTag fusion, (ii) broadening the range of effective photosensitizers, and (iii) optimizing irradiation parameters to enable precise spatial control over filament disassembly. Together, these findings position FilaBuster as a robust platform for acute, selective, and spatiotemporally precise disassembly of IF networks, enabling new investigations into their structural and functional roles in cell physiology and disease.

## Introduction

Intermediate filaments (IFs) are a diverse family of apolar, semiflexible cytoskeletal fibers that maintain nuclear shape, support organelle positioning, and reinforce tissue integrity. Over 70 IF genes have been identified in metazoans, spanning multiple subtypes—including keratins, vimentin, desmin, neurofilaments, and nuclear lamins—with distinct structural and tissue-specific roles. Mutations or dysregulation in IFs are implicated in a wide range of diseases, from cardiomyopathies and neurodegeneration to cancer.^1,2^

Despite their biological and clinical relevance, IFs remain uniquely challenging to perturb with both speed and specificity. While actin filaments and microtubules can be acutely disassembled using well-established small-molecule inhibitors,^3,4^ IFs lack fast-acting, selective tools. Existing IF disrupting agents—including phosphatase inhibitors,^5^ steroidal lactones,^6^ oxidizing agents,^7^ aliphatic alcohols,^8,9^ and electrophilic toxins^10,11^—typically induce broad cellular stress and off-target effects,^12^ limiting their interpretability. Gene silencing or knockout methods offer more precision, but require days to manifest and often trigger compensatory responses that obscure acute IF functions. As a result, key questions about IF architecture, dynamics, and mechanobiology remain unresolved.

To address this methodological gap, we investigated whether the inherent sensitivity of IFs to oxidative stress^13–15^ could be leveraged to create a selective and controllable IF disassembly strategy. Rather than applying high concentrations of diffusible oxidants to the entire cell, we sought to generate reactive oxygen species (ROS) in situ, directly and selectively at the filament surface. Photosensitizers—molecules that produce ROS upon illumination—offer a compelling mechanism for achieving this. Their activity is tightly constrained in both space and time: the reactive intermediates they produce act over nanometer scales^16^ and persist for only microseconds.^17^ Photosensitizers have been used in chromophore-assisted light inactivation (CALI) to acutely inhibit tagged proteins,^18–20^ as well as to damage organelles such as mitochondria^21^ under intense, localized irradiation. We reasoned that if photosensitizers could be densely and specifically distributed throughout the IF network, their activation would produce highly localized oxidative stress sufficient to trigger filament disassembly—without requiring global ROS elevation or widespread cellular damage.

To implement this concept, we developed FilaBuster, a modular photochemical strategy that enables rapid, specific, and tunable IF disassembly. The strategy localizes ROS generation directly at the filament surface using a three-part framework (Fig. 1a). First, an IF-HaloTag fusion protein (e.g., GFAP-HaloTag) is ectopically expressed and co-assembles with native filaments, embedding the HaloTag domain stably within the endogenous IF network. Second, the HaloTags are covalently labeled with cell-permeable, photosensitizer dyes (e.g., Janelia Fluor 570 HaloTag ligand, JF_570_-HTL),^22^ thereby arming the system for light-induced activation. Third, illumination triggers ROS production from the bound photosensitizers, leading to spatially restricted oxidative damage and rapid filament fragmentation. Here, we describe the development and validation of this approach, demonstrating its speed, specificity, and versatility across multiple IF systems.

**Figure 1:**
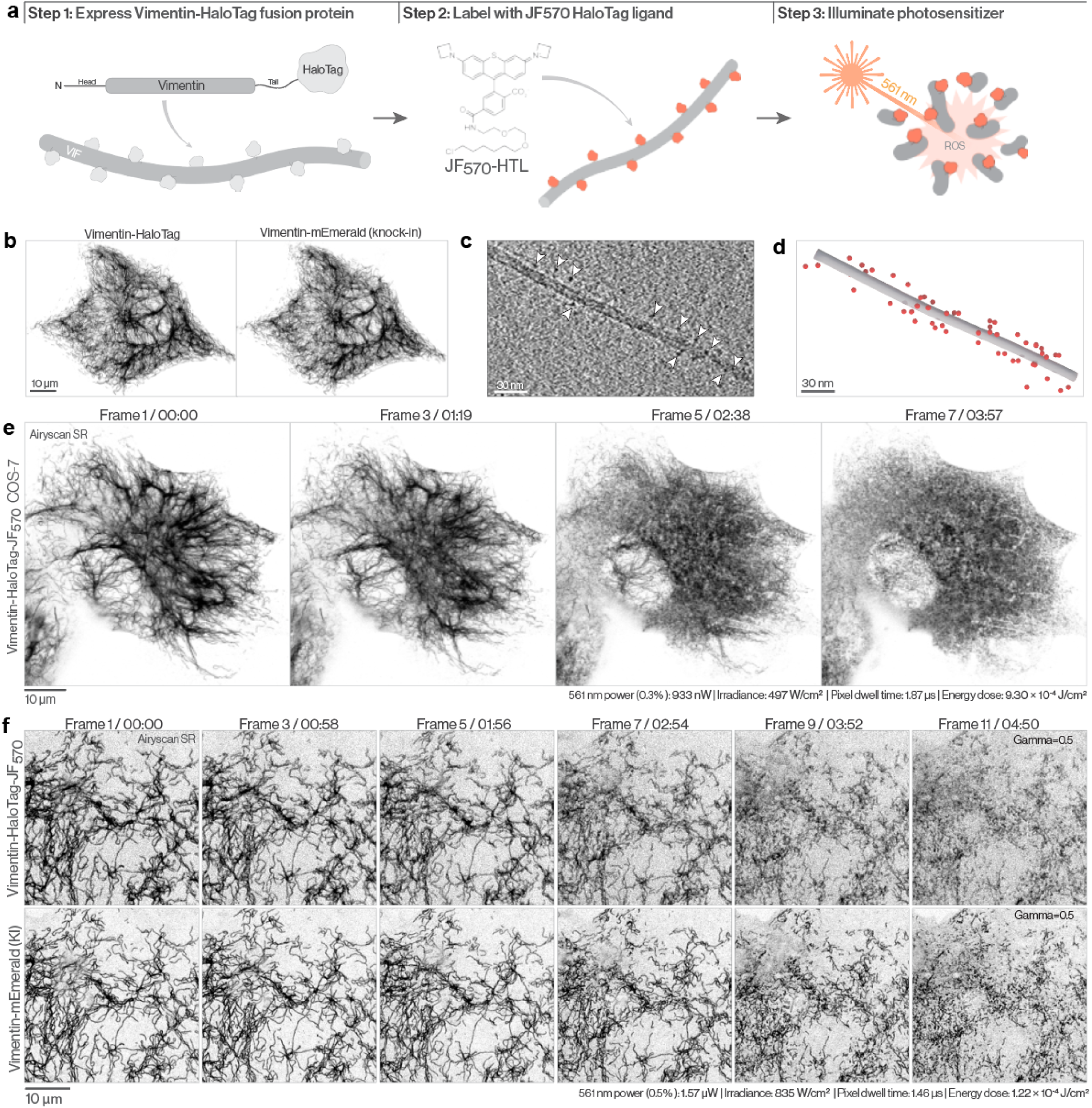
Acute fragmentation of vimentin IFs upon excitation of a vimentin-targeted photosensitizer. **a.** Schematic outlining the FilaBuster implementation. Step 1: Vimentin-HaloTag is expressed and co-assembles with native filaments. Step 2: Cells are labeled with JF_570_-HTL, a cell-permeable photosensitizer that covalently binds HaloTag. Step 3: 561 nm light excites the photosensitizer dye, generating ROS to drive localized filament disassembly. **b.** Airyscan images showing colocalization of stably expressed vimentin-HaloTag (JFX_549_, left) with endogenously tagged vimentin-mEmerald (right) in COS-7 cells. **c.** Tomogram slice of a VIF from a vimentin-HaloTag-expressing MEF, with HaloTag densities indicated by white arrows. **d.** 3D model of a VIF (gray) with associated HaloTags (red spheres) reconstructed from cryo-ET. **e.** Airyscan montage of a COS-7 cell expressing vimentin-HaloTag labeled with JF_570_-HTL. 561 nm illumination induces progressive VIF disassembly over ∼4 minutes. **f.** Airyscan montage of endogenously tagged vimentin-mEmerald (top row) and co-expressed vimentin-HaloTag-JF_570_ (bottom row) in a COS-7 cell (images in f are gamma adjusted (0.5) for improved visualization of filaments). 561 nm laser power (measured at the sample plane), irradiance, pixel dwell time, and energy dose per pixel are indicated for panels e and f.

## Results

### Vimentin-HaloTag incorporates into native vimentin IFs and meets spatial requirements for targeted oxidative disassembly

To prototype the FilaBuster approach, we initially focused on vimentin IFs as a proof-of-concept model system. Vimentin is among the best-characterized IF proteins,^23^ forms simple homopolymers^24^, and its complete filament structure has recently been resolved,^25^ making it an ideal starting point for validating and refining the FilaBuster strategy before extending it to other IF systems. We designed a fusion protein linking HaloTag7^26^ to the C-terminal tail of human vimentin. To establish its specificity and proper integration into the native vimentin IF (VIF) network, we first stably expressed vimentin-HaloTag in wild-type COS-7 cells. After fixation and immunostaining, Airyscan confocal microscopy revealed extensive colocalization between the HaloTag fusion and immunolabeled filaments, confirming proper incorporation into endogenous VIFs (Fig. S1a-b). To further validate native VIF integration in live cells, we expressed the construct in vimentin-mEmerald knock-in COS-7 cells, observing complete overlap between vimentin-HaloTag and endogenously-tagged vimentin (Fig. 1b).

Fluorescence microscopy confirmed the specificity of our fusion protein, but to determine whether its incorporation affected the structural integrity of individual VIFs, we applied cryo-electron tomography (cryo-ET) to permeabilized, plunge-frozen mouse embryonic fibroblasts^27^ (MEFs) stably expressing vimentin-HaloTag. Tomogram analysis revealed intact vimentin filaments, approximately 11 nm in diameter, surrounded by 3–4 nm spherical densities, consistent with the dimensions of HaloTag (Fig. 1c-d; Fig. S1c, Video 1). HaloTags were positioned at a mean distance of 12.27 ± 3.75 nm from the filament center (Fig. S1d), with an average of 1.94 ± 0.43 HaloTag proteins per 10 nm of filament length. Given that each 10 nm segment of vimentin contains 12–14 tail domains,^25^ this corresponds to a mean HaloTag occupancy of ∼15% (Fig. S1e). Together, these findings indicate that the vimentin-HaloTag fusion integrates into endogenous VIFs without compromising filament architecture. Furthermore, the HaloTag proteins are positioned in close proximity to the filament surface—within a range that permits effective interaction with short-lived ROS generated by photosensitizing ligands. Finally, the uniform distribution of HaloTags along the filament ensures extensive coverage, enhancing the likelihood of comprehensive and efficient filament disassembly upon photosensitizer labeling and subsequent irradiation.

### Controlled disassembly of VIFs by activation of a HaloTag-targeted photosensitizer

Having established that our probe effectively integrates into endogenous filaments, we next assessed its functional utility for light-mediated VIF disassembly. To this end, we labeled vimentin-HaloTag expressing cells with the photosensitizer JF_570_-HTL (250 nM, 30 min) and attempted to induce VIF disruption using 561 nm illumination. To prevent premature activation, cells were initially monitored under transmitted light and appeared healthy with normal morphology. We then acquired Airyscan (Fig. 1e) or widefield (Fig. S2a) time-lapse movies using low-power 561 nm laser illumination (see methods/figure legends for details). In the initial frames, VIFs displayed normal morphology, forming an extensive filamentous network extending throughout the cytoplasm. However, beginning at approximately five illumination cycles, we observed progressive and widespread VIF fragmentation, accompanied by a steady rise in diffuse cytoplasmic fluorescence consistent with increasing levels of soluble vimentin subunits (Fig. 1e, Fig. S2a, Video 2).

To quantitatively assess VIF disassembly, we evaluated three distinct metrics that reflect filament network integrity. First, the coefficient of variation—a measure of fluorescence intensity heterogeneity—decreased over time, consistent with the disappearance of dense filament bundles and a shift toward a more uniform, diffuse signal (Fig. S2b). Second, the radial intensity profile, which reflects the spatial distribution of VIFs from nucleus to periphery, progressively flattened, consistent with the dispersal of VIF fragments away from the perinuclear region (Fig. S2c). Finally, local image coherency, a measure of directional filament alignment, dropped sharply, reflecting the breakdown of elongated filaments into isotropic fragments (Fig. S2d).

To confirm that our approach disassembles native vimentin filaments, we repeated the experiment in vimentin-mEmerald knock-in COS-7 cells co-expressing vimentin-HaloTag-JF_570_. In these cells, we observed concurrent fragmentation of both HaloTag- and mEmerald-labeled VIFs, confirming that light-induced disruption extends to the endogenous filament network (Fig. 1f).

### Photochemical disassembly of native VIFs via nanobody-HaloTag fusions

We found that the vimentin-HaloTag fusion seamlessly integrates into native VIFs, providing an effective means of stably anchoring photosensitizer ligands at the filament surface. However, introduction of a tagged IF subunit requires careful validation to ensure that the fusion does not interfere with normal filament dynamics.^28^ Moreover, ectopic expression of IF subunits - even untagged - can increase the density of the IF network and meaningfully alter its mechanical properties.^23^ While these concerns can be mitigated with careful controls and thoughtful experimental design, genetic tagging of IF subunits may not always be feasible or desirable.

An attractive alternative to direct subunit tagging is the use of live cell expressible nanobody directed against IF proteins.^29^ Comprising a conventional fluorescent protein fused to the variable domain (V_H_H) of an Alpaca heavy chain antibody, fluorescent nanobodies, or chromobodies, have been developed to label multiple IF subtypes, including keratin 8, lamin, and vimentin.^30^ We hypothesized that we could use a nanobody-HaloTag fusion to direct photosensitizer ligands to native IFs in live cells, effectively circumventing concerns about direct subunit tagging or overexpression.

To explore the viability of this approach, we generated a construct linking HaloTag to the C-terminus of a well characterized vimentin nanobody^31^, and expressed the fusion (vimentin-V_H_H-HaloTag) in vimentin-mEmerald knock in COS-7 cells (Fig. S3a). Fluorescence microscopy confirmed specific binding and localization of the nanobody fusion to the native vimentin network without altering its organization. Labeling vimentin-V_H_H-HaloTag expressing cells with JF_570_-HTL followed by 561 nm illumination induced rapid disintegration of endogenous vimentin filaments, with comparable efficiency to what we had observed with vimentin-HaloTag-JF_570_ (Fig. S3b-d, Video 3). This nanobody-based targeting strategy thus significantly broadens the scope and applicability of FilaBuster, enabling targeted, photochemical disassembly of VIFs without the requirement for direct vimentin tagging. Given the availability of validated nanobodies against other IF proteins,^30^ this strategy should be readily adaptable to target additional filament classes.

### Expanding FilaBuster with a red-shifted, fluorogenic HaloTag photosensitizer for controlled VIF disassembly

Building upon the successful induction of rapid VIF disassembly with JF_570_, we aimed to broaden the utility of FilaBuster by developing a red-shifted HaloTag photosensitizer ligand that could be activated using far-red lasers common on standard fluorescence microscopes. Such a ligand would facilitate deeper tissue penetration due to the longer excitation wavelength and enable multiplexed imaging with conventional blue and green-excited fluorophores, further enhancing the versatility of our approach.

To this end, we synthesized JF_634_, a derivative of JF_608_ modified with bromine substituents to enhance intersystem crossing efficiency via the heavy-atom effect (Fig. S4a). To allow covalent labeling of HaloTag fusion proteins, we further functionalized JF_634_ with a chloroalkane linker, generating JF_634_-HTL. Spectroscopic characterization of JF_634_-HTL in the presence of HaloTag enzyme revealed a peak absorbance at 639 nm as well as a ∼300-fold increase in absorbance compared to the unbound ligand (Fig. S4a-c).

We next evaluated JF_634_-HTL in live cells stably expressing vimentin-HaloTag (Fig. S4d). The dye exhibited efficient cellular uptake and selective labeling of the filament network, with no detectable cytotoxicity or vimentin cytoskeletal artifacts following prolonged incubation in the dark. Upon illumination with 639 nm (Airyscan) or 640 nm (widefield) light, vimentin-HaloTag–JF_634_ triggered robust and rapid VIF disassembly in both COS-7 and MEF (Fig. S4e–f, Video 4), with disassembly kinetics comparable to, and in some cases faster than, those observed with JF_570_. Together, these results establish JF_634_-HTL as an effective, red-shifted alternative to JF_570_.

### Dose-dependent control of VIF fragmentation using conventional HaloTag ligands

While the two validated vimentin-targeted photosensitizers induce rapid VIF fragmentation upon low-light minimal illumination conditions, their extreme sensitivity to excitation light precludes extended imaging of native VIF dynamics prior to targeted disruption. To overcome this limitation, we sought dyes capable of operating in two distinct modes: a low-power “standard imaging” mode for visualizing intact IFs, and a high-power “FilaBuster” mode for inducing filament fragmentation (Fig. 2a).

**Figure 2:**
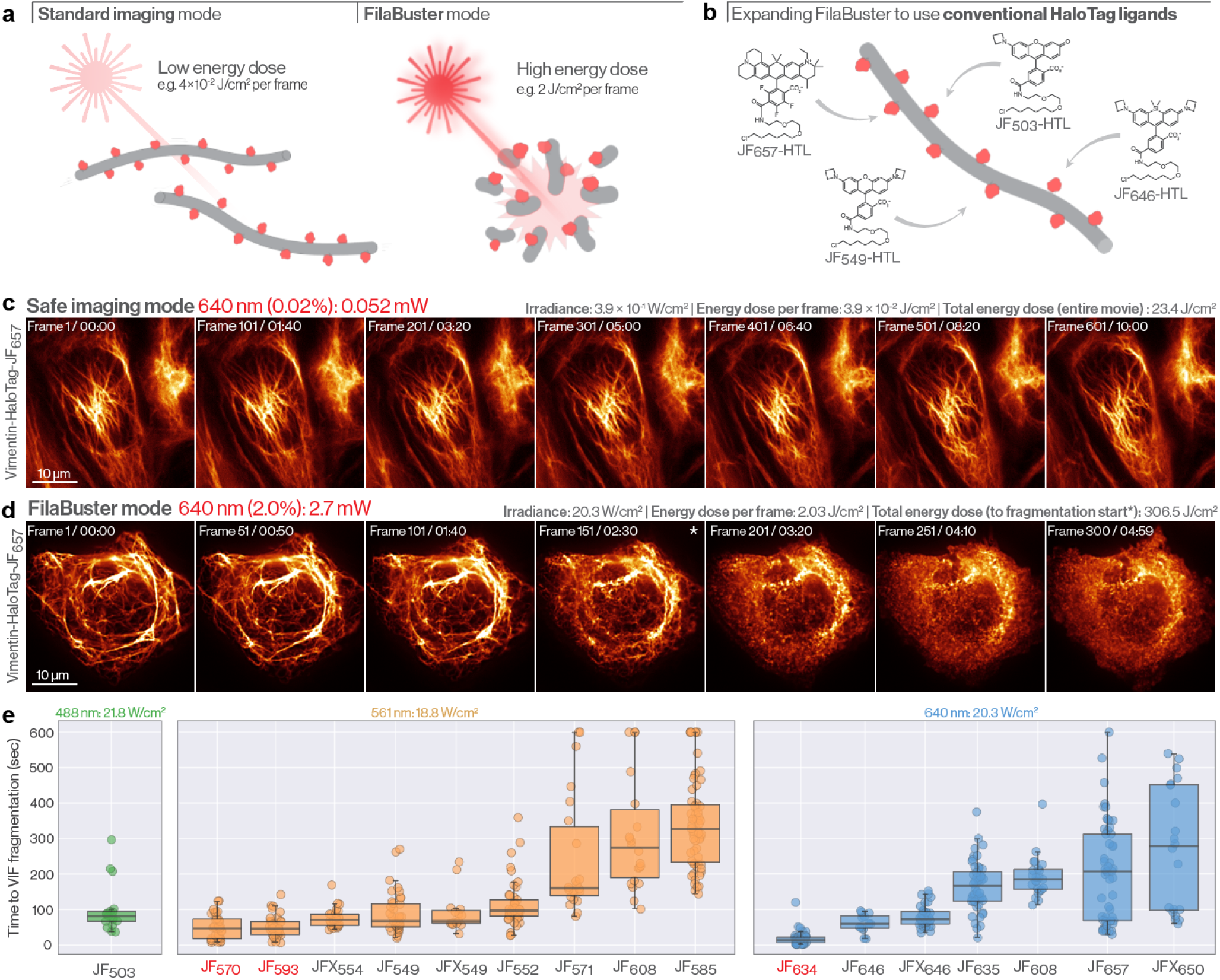
Expanding the FilaBuster strategy with conventional HaloTag ligands. **a.** Schematic indicating the effect of light-dose on conventional HTLs targeted to VIFs. In low-power “standard imaging mode,” vimentin-HaloTag labeled with a conventional HTL can be imaged for extended periods without disrupting filament integrity. In “FilaBuster mode,” enhanced illumination results in VIF fragmentation. **b.** Cartoon depicting various conventional, non-photosensitizer ligands that can be used to label VIF-targeted HaloTag. **c.** Widefield epifluorescence timelapse of a MEF expressing vimentin-HaloTag-JF_657_ acquired in standard imaging mode. Over 601 illumination cycles with low 640 nm laser power (0.052 mW) we observe no detectable VIF fragmentation. **d.** Timelapse of a MEF expressing Vimentin-HaloTag-JF_657_ acquired in “FilaBuster mode.” Illuminating with ∼ 50× higher 640 nm laser power, we observe clear VIF fragmentation within ∼150 cycles. **e**. Average duration to VIF fragmentation in vimentin-HaloTag MEFs labeled with the indicated HaloTag ligand (red=photosensitizer, gray=conventional ligand). Cells were excited at either 488 nm (green plot), 561 nm (orange plots), or 640 nm (blue plots) at 1 frame per second, 100 ms exposure, and the indicated irradiance. Time to VIF fragmentation was recorded for each dye. JF_608_ was screened with both 561 nm and 640 nm excitation. Cells in which VIF fragmentation failed to occur within 10 minutes were plotted at 600 seconds. 640 nm laser power (measured at the sample plane), irradiance, exposure time, energy dose per frame, and total energy dose are indicated for panels c and d.

We hypothesized that conventional rhodamine-based HaloTag ligands, despite lacking heavy-atom substituents for efficient singlet oxygen production, might exhibit latent photoreactivity under elevated illumination intensities (Fig. 2b). To test this, we labeled vimentin-HaloTag MEFs with JF_657_-HTL, a bright, far-red–excitable carborhodamine, and imaged cells with a 640 nm laser in either low-power standard mode (irradiance: 0.39 W/cm²) or high-power FilaBuster mode (irradiance: 20.3 W/cm²). In standard mode, VIFs remained intact and morphologically stable over ten minutes or 601 cycles of illumination (Fig. 2c). In contrast, under FilaBuster mode, robust filament disassembly was triggered after ∼150 frames (Fig. 2d), requiring a cumulative energy dose of 307 J/cm²—over 13-fold greater than the total energy dose over 10 minutes in standard imaging mode (Fig. 2c-d). These findings demonstrate that conventional HTLs can reliably trigger VIF disassembly when activated at sufficiently high energy levels, despite their lower intrinsic photoreactivity.

### Benchmarking conventional HTLs as FilaBusters

Encouraged by these results, we systematically benchmarked 13 conventional HTLs^32–36^ and 3 photosensitizer HTLs^37^ in vimentin-HaloTag MEFs using widefield epifluorescence microscopy under FilaBuster-mode imaging conditions (Fig. S5a). Surprisingly, every ligand successfully induced VIF fragmentation, though with varying efficiency (Fig. 2e, Fig. S5b-c, Video 5). As predicted, photosensitizers such as JF_634_-HTL were the most potent, triggering network collapse within ∼17 cycles of 640 nm irradiation. However, several conventional HTLs, including JF_549_, JF_646_, and their deuterated JFX analogs, were also highly effective, driving filament disassembly within 60–90 higher-energy illumination cycles. Illumination parameters and corresponding energy thresholds for each ligand are summarized in Fig. S5b. While variation in ligand performance likely reflects differences in intersystem crossing efficiencies, other factors such as dye loading and excitation efficiency at the applied laser line may also contribute (Fig. S5c).

The observation that all tested ligands were capable of driving VIF disassembly under elevated illumination underscores the broad compatibility of rhodamine-based HTLs with the FilaBuster framework. Moreover, we observed similarly efficient VIF disassembly across a wide range of cell types, including epithelial lines, immortalized fibroblasts, and primary human fibroblasts (Fig. S6), demonstrating that FilaBuster performance generalizes across diverse cellular contexts.

### Operational boundaries of FilaBuster-induced IF disassembly

Encouraged by the broad efficacy of conventional and photosensitizer HaloTag ligands in driving IF disassembly, we next sought to define the mechanistic constraints that govern FilaBuster’s activity. Specifically, we asked how tightly the system depends on spatial targeting, molecular configuration, and the redox sensitivity of the underlying filament network.

To test the spatial proximity requirements of FilaBuster, we asked whether ROS generation must occur directly at the filament surface or whether non-targeted diffusely distributed photosensitizers throughout the cytoplasm could still drive light-mediated VIF disassembly. As a test of this boundary condition, we expressed an unfused, cytosolic HaloTag, labeled it with JFX_549_-HTL, and subjected cells to high-intensity 561 nm laser illumination (Fig. S7a-b). Despite robust ligand photobleaching and energy doses that far exceeded those required to fragment vimentin filaments labeled with the same dye, no VIF disruption was observed (Fig. S7b). These findings reveal a strict spatial requirement for photosensitizer activity: ROS must be generated at the filament surface to induce IF disassembly, consistent with their nanometer-scale diffusion ranges and rapid quenching in the intracellular environment.

We then asked whether other intracellular targets are similarly susceptible to localized ROS when delivered via the FilaBuster targeting system. Specifically, we tested whether directing JF_634_ to F-actin via a LifeAct–HaloTag fusion would result in actin filament fragmentation under illumination conditions able to induce VIF fragmentation (Fig. S7c). We irradiated cells with 633 nm light (peak irradiance: 4012 W/cm^2^), but observed no obvious actin filament disassembly or off-target effects on VIFs labeled with a conventional vimentin-mEmerald reporter (Fig. S7c, Video 6). These observations indicate that FilaBuster-induced disruption is not a generalizable effect of local ROS production, but a consequence of targeting a uniquely vulnerable cytoskeletal substrate. Importantly, the integrated energy doses used to drive FilaBuster-mediated VIF disassembly—as little as 34.5 J/cm² in the case of vimentin-HaloTag-JF_634_—are one to three orders of magnitude lower than those required to induce comparable effects in classical CALI systems,^19,22,38–41^ which often exceed 10,000 J/cm² (Fig. S7d).

To test whether the photochemical mechanism requires the specific use of HaloTag and chloroalkane ligands, we evaluated two alternative strategies for localizing ROS generation to the filament surface. First, we generated a vimentin–SuperNova2^42^ fusion to produce VIF localized ROS via a genetically encoded fluorescent photosensitizer. Upon prolonged 561 nm illumination we observed complete photobleaching of SuperNova2, but no observable VIF disassembly assessed using a co-expressed vimentin-mEmerald marker (Fig. S8a, Video 6). Failure of the vimentin–SuperNova2 approach likely reflects two inherent limitations of fluorescent proteins compared to small-molecule dyes: substantially lower singlet oxygen and superoxide quantum yields^43^ and chromophores that are shielded by the β-barrel structure, potentially limiting ROS diffusion to nearby filament targets. Second, we labeled vimentin–SNAP-tag fusions with the JFX_554_ SNAP-tag ligand and subjected cells to high-power 561 nm illumination. Here, we observed minimal and delayed fragmentation, only after extended exposure and substantial photobleaching (Fig S8b, Video 6). The reduced efficiency relative to HaloTag may reflect slower ligand binding kinetics^44^, electrostatic differences between tags (pI 9.6 for SNAP vs. 4.9 for HaloTag), or less favorable spatial positioning of the dye relative to the filament surface.

Taken together, these results define the spatial, structural, and mechanistic boundaries of the FilaBuster system. Efficient disassembly requires not only ROS production, but also precise positioning of the photosensitizer at native IFs. The failure of alternative tag systems and the resistance of unrelated cytoskeletal structures to equivalent light doses underscore that FilaBuster is not a generic photosensitizer platform, but a designed system for targeted, efficient, and selective fragmentation of inherently ROS-sensitive IFs.

### Light-Induced VIF fragmentation is associated with increased vimentin methionine oxidation

Given the well-established sensitivity of vimentin to oxidative stress,^10^ and extensive evidence implicating its single cysteine residue (C328) in redox-mediated filament disassembly,^45^ we initially hypothesized that FilaBuster operates by targeting this residue. However, C328 proved dispensable. Vimentin–HaloTag constructs harboring either a C328A or C328H mutation disassembled upon photosensitizer activation with kinetics indistinguishable from wild-type. This held true both in cells with an untagged endogenous vimentin background and in vimentin-null MEFs reconstituted exclusively with mutant constructs, confirming that filament fragmentation does not require modification at C328 (Fig. S9).

Having ruled out cysteine as the critical target, we considered whether direct photochemical crosslinking between dye and filament might explain the effect since a light-induced reaction between BSA and rhodamine dyes containing cyclic amine substituents has been observed in vitro^46^. However, we observed robust VIF fragmentation with rhodamine dyes containing acyclic amine substituents, such as SiTMR, as well as with the carbofluorescein (CFl) HaloTag ligand, which lacks alkylamines entirely, arguing against covalent adduct formation as a general mechanism.

To directly assess oxidative modifications, we performed mass spectrometry on vimentin–HaloTag U2-OS cells processed under three conditions: dye labeling + irradiation, dye labeling without irradiation, and unlabeled controls. Quantitative analysis revealed a striking increase in oxidized methionine residues in the irradiated group across multiple sites (Fig. S10a). This pattern is consistent with targeted ROS generation at the filament surface and suggests that localized subunit oxidation likely underlies disassembly. Of note, fluorescent detection of cellular ROS using CellROX Deep Red showed no measurable increase in signal following FilaBuster activation (Fig S10b), suggesting minimal oxidative spillover and reinforcing the idea that disassembly is driven by highly localized photochemical events.

Together, these results indicate that FilaBuster operates by generating ROS at the filament surface, which in turn oxidizes vimentin subunits—primarily at methionine residues—triggering filament destabilization. This mechanism is not dependent on C328, does not involve bulk oxidative stress outside of the target filament, and can be tuned by varying irradiation parameters and HaloTag ligand identity.

### FilaBuster enables spatially controlled VIF fragmentation

We reasoned that by confining high-energy “FilaBuster” illumination to defined subcellular regions, we could induce localized fragmentation of the VIF network while preserving surrounding filaments (Fig. 3a), enabling spatiotemporally resolved perturbation of IF architecture—an approach that, until now, has not been possible.

**Figure 3:**
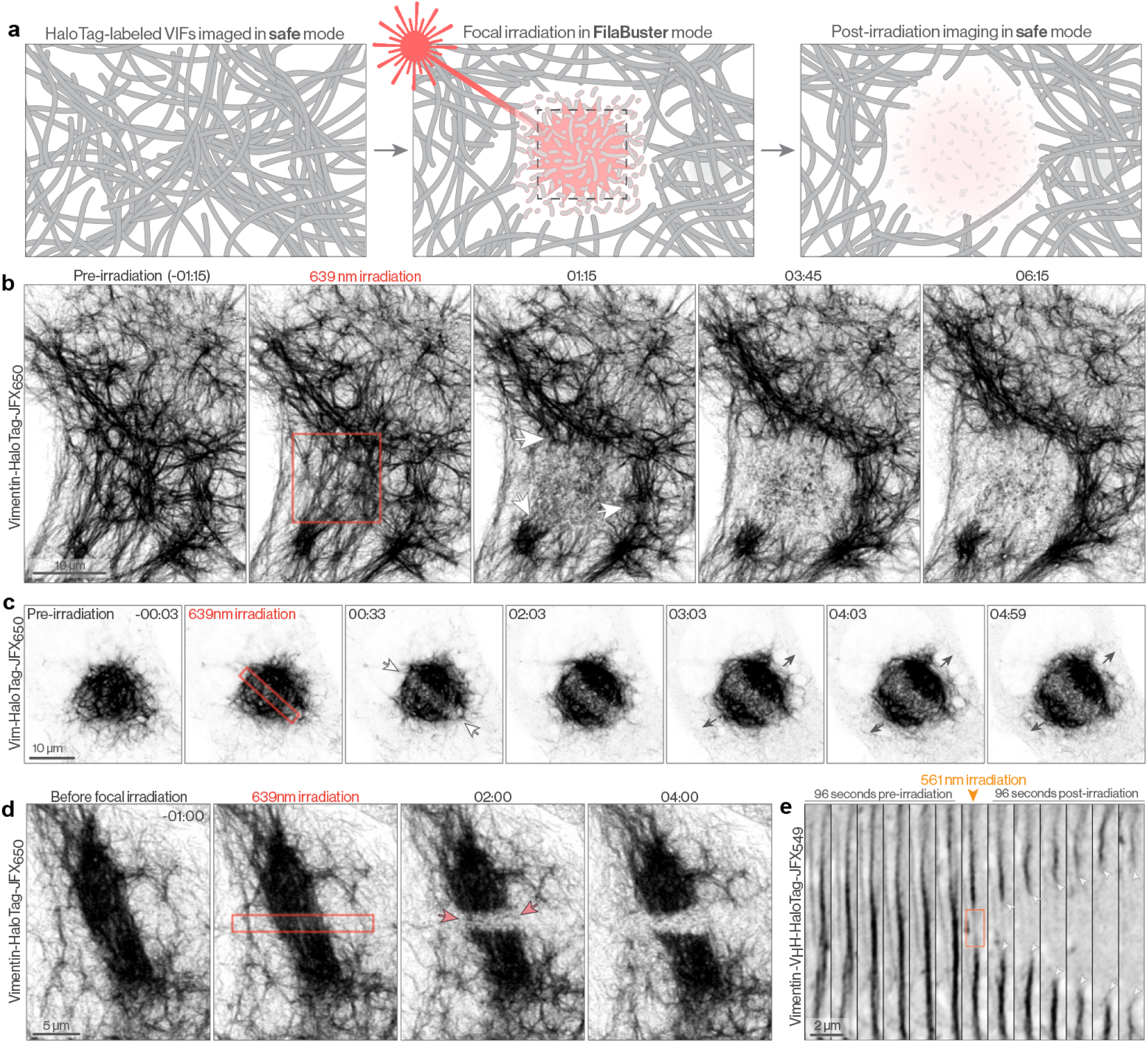
FilaBuster enables precise spatial control over VIF fragmentation. **a.** Cartoon schematic indicating focal irradiation strategy for targeted VIF disassembly. **b.** Airyscan montage of a COS-7 stably expressing Vimentin-HaloTag-JFX_650_. The cell was focally irradiated with a 639 nm laser at the indicated region (red box), resulting in localized VIF disassembly. **c.** Focal irradiation of a dense, perinuclear vimentin cap. Within 30 seconds of 639 nm irradiation, VIFs within the irradiation window (red box) fragment and the cap is bisected. Over several minutes, the two hemispheres of the cap continue to separate, and we observe a subtle increase in soluble vimentin background in the cytoplasm. **d.** Airyscan montage of a dense vimentin IF bundle labeled with vimentin-HaloTag-JFX_650_. Brief 639 nm irradiation at the indicated region (red box) resulted in targeted IF disassembly, effectively bisecting the bundle within 2 minutes. **e.** Airyscan montage of an extended vimentin IF bundle labeled with vimentin-V_H_H-HaloTag-JFX_549_. Focal 561 nm irradiation (red box) severs the bundle, resulting in retraction of the two free ends (arrows).

To test this, we performed time-lapse imaging of COS-7 cells stably expressing vimentin-HaloTag and labeled with JFX_650_-HTL, a rhodamine-based dye with low photochemical reactivity under standard imaging conditions. Cells were initially imaged under low-power standard conditions to establish a stable baseline. At a defined time point, a ∼10 × 10 µm region of interest (ROI) was exposed to focused 639 nm illumination at higher irradiance to trigger FilaBuster mode activation. Imaging then resumed under standard, low-power imaging conditions to monitor structural changes post-irradiation. Within one minute, vimentin IFs within the irradiated ROI underwent rapid and complete disassembly, while filament structures outside the targeted area remained intact (Fig. 3b, Video 7). Notably, we did not observe VIF transport into the irradiated region as typically observed when photobleaching conventional vimentin fusion proteins (Fig. S11a), suggesting that the irradiated VIF breakdown products either immediately destabilize any newly trafficked VIFs or interfere with local VIF trafficking.

Furthermore, focal illumination at wavelengths outside the FilaBuster ligand’s excitation spectrum did not induce VIF disintegration. In cells co-expressing vimentin-mNeonGreen and Vimentin-HaloTag-JFX_646_, 488 nm irradiation had no effect on VIF stability while 639 nm irradiation drove robust fragmentation (Fig. S11b-c).

Building on this spatial precision, we next explored whether we could disrupt structurally diverse vimentin IF assemblies. By positioning focal irradiation windows over thick, bundled IF structures, we demonstrated that brief and spatially confined 639 nm irradiation efficiently bisected perinuclear vimentin caps as well as thick peripheral VIF bundles (Fig. 3c-d).

Focal VIF fragmentation could also be achieved using the nanobody targeting strategy (Video 8). With vimentin-V_H_H-HaloTag-JFX_549_, we were able to precisely sever thin filament bundles with focal 561 nm illumination, leading to dramatic retraction of the free ends (Fig. 3e). To ensure that disassembly could be robustly visualized with minimal interference from imaging conditions, we repeated the experiment using a co-expressed vimentin-mEmerald reporter (Fig. S11d, Video 8). This approach allowed us to monitor the same irradiation-induced disassembly process while exclusively using the stable mEmerald signal for before-and-after imaging. Since mEmerald is excited with the 488 nm laser line, which does not activate the photosensitizer, this method ensured that IF disassembly was induced solely by targeted irradiation and not by imaging-associated illumination.

All validated HaloTag ligands from our global disassembly assays (Fig. 2e) supported localized filament disruption under focal irradiation. However, potent photosensitizer dyes like JF_570_-HTL posed challenges for spatial precision due to their high reactivity; even minimal light exposure after focal irradiation caused widespread filament loss (Movie 6).

### Rapid, localized filament scission defines early-stage VIF disassembly kinetics

To define the kinetics and mechanical characteristics of FilaBuster-mediated disassembly, we performed high-speed TIRF and confocal imaging of labeled vimentin filaments before, during, and after illumination-induced fragmentation. Under continuous illumination, filaments initially appeared stable and phenotypically unremarkable. However, upon exceeding the threshold light dose, we observed the sudden onset of filament disassembly, typically initiated by 2-3 severing events, producing ∼1 µm truncated filaments. Severing frequently occurred at points of high curvature, as seen in both TIRF and Airyscan recordings (Fig. 4a–b). Truncated filaments persisted briefly before collapsing into minimal, insoluble fragments (Fig. 4c).

**Figure 4:**
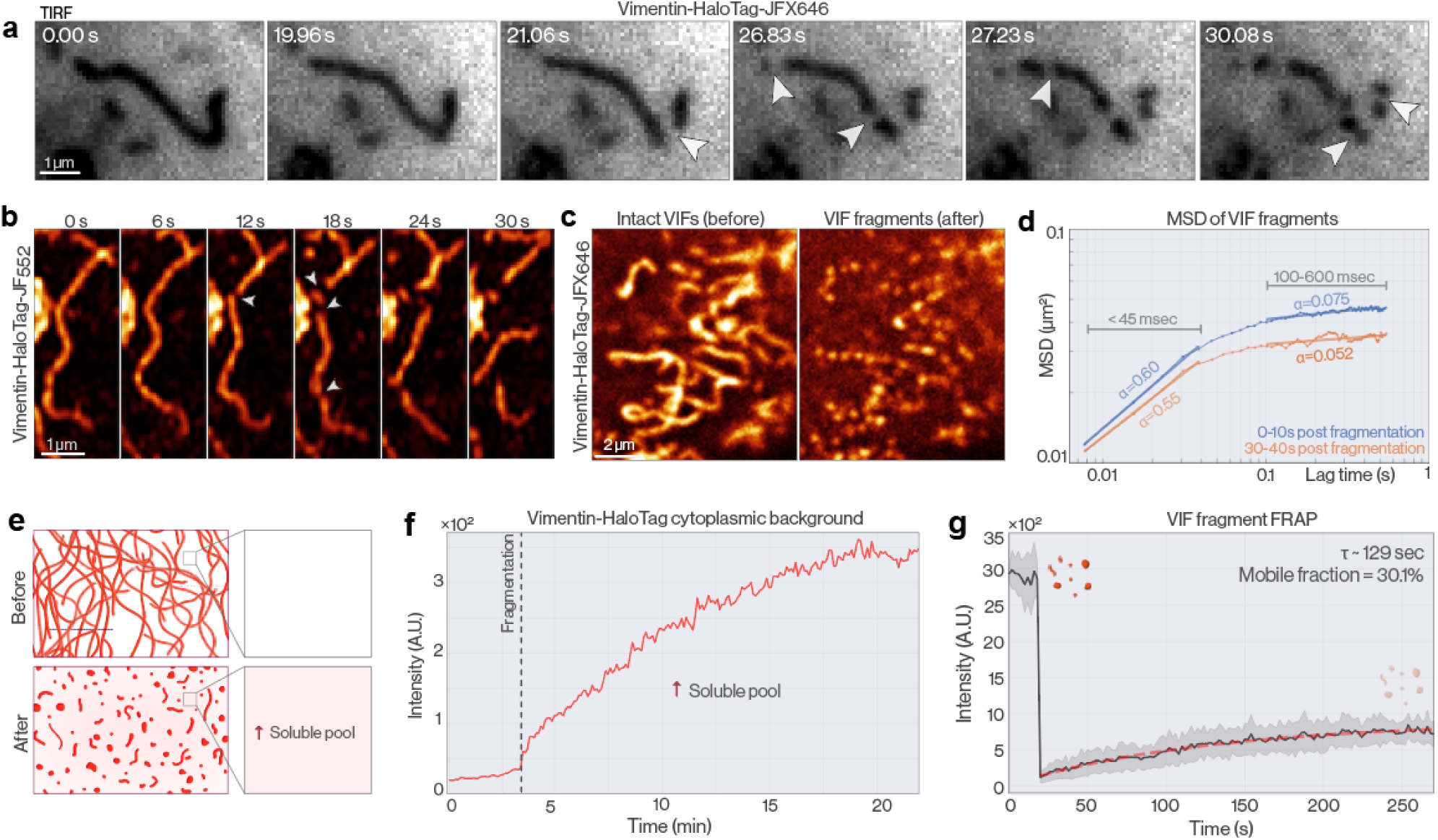
Kinetics of VIF fragmentation. **a**. TIRF montage of VIF fragmentation in a U2-OS cell expressing vimentin-HaloTag-JFX_646_. Arrows indicate sites of filament scission. **b.** Airyscan timelapse capturing fragmentation of a single VIF labeled with vimentin-HaloTag-JF_552_ in a COS-7 cell. **c.** Comparison of VIFs before (left) or immediately after (right) fragmentation. **d.** MSD of VIF fragments computed from single particle tracking of VIF fragments either in the first 10 seconds after VIF disassembly (blue), or 30-40 seconds after VIF disassembly (red). In each case, alpha values were computed over short (<45 msec) and longer (100-600 msec) lag times. **e.** Cartoon schematic indicating VIF fragmentation and increase in soluble VIF subunits using the FilaBuster approach. **f** Global VIF fragmentation is accompanied by a rapid increase in diffuse vimentin-HaloTag signal in regions devoid of filaments prior to irradiation, consistent with increased soluble VIF subunits. **g**. Fluorescence recovery after photobleaching (FRAP) of confined VIF fragments display modest recovery over 4 minutes, consistent with continued subunit exchange.

To characterize the behavior of newly formed VIF fragments, we performed single-particle tracking during the first few seconds post-fragmentation. (Fig 4d). Mean-squared displacement (MSD) analysis revealed a two-phase motility profile. At short lag times (<45 ms), fragments exhibited subdiffusive motion (α ≈ 0.60, D_eff_ ≈ 0.22 µm^2^/s), consistent with constrained diffusion through the cytoplasm. At longer lag times (100–600 ms), the MSD plateaus (α ≈ 0.075). We interpret this motility state transition as a consequence of polydisperse VIF fragments attaching and detaching with the cytoplasmic meshwork of actin filaments and microtubules^47^. Concurrent with VIF fragmentation, we observed a global rise in diffuse, cytoplasmic vimentin-HaloTag signal, even in regions previously devoid of assembled VIFs; an effect we attribute to a dramatic increase in the typically negligible soluble vimentin tetramer pool (Fig. 4e-f). Together, these data suggest that FilaBuster-induced disassembly generates at least two distinct end-products: spatially confined, truncated VIF fragments and a rapidly diffusing soluble vimentin pool—each with potentially distinct cellular effects.

### VIF fragments persist in the cytoplasm and block new filament assembly

While the early phase of disassembly results in widespread filament fragmentation and redistribution of soluble subunits, we next asked whether the network could reassemble after this acute disruption. To determine whether VIF fragments retained dynamic subunit exchange with the soluble pool, we performed fluorescence recovery after photobleaching (FRAP) on stationary fragments ∼10 minutes after disassembly (Fig 4g). Over 250 seconds, we observed ∼30% recovery with a characteristic time (τ) of ∼129 seconds, indicating that fragments retain the capacity for subunit exchange, but not for productive reassembly. Instead, over the subsequent hours, these fragments gradually coalesced, migrating toward the perinuclear region at ∼25 nm/min and condensing into dense, immobile aggregates ranging from 500 nm to 5 µm in diameter (Fig. S12a–b).

To test whether this assembly block extended to newly synthesized vimentin, we performed a two-color pulse-chase assay. Cells were first labeled with JF_549_ to mark the pre-existing vimentin-HaloTag pool, then immediately subjected to 561 nm light-induced disassembly. After a 20-hour recovery period, we labeled the cells with JF_635_-HTL to visualize any vimentin-HaloTag proteins synthesized after irradiation. If filament assembly were possible, we would expect these newly translated, non-irradiated subunits to form a distinct filamentous network. Instead, JF_635_-labeled vimentin failed to assemble and co-localized with the original JF_549_-labeled aggregates, despite never being exposed to light (Fig. S12c). Together, these results demonstrate that FilaBuster not only irreversibly fragments irradiated VIFs, but also imposes a dominant, long-lasting block on reestablishing a filamentous network, even in the presence of abundant, newly synthesized vimentin.

### Acute VIF fragmentation reveals immediate effects on cell morphology and cytoskeletal organization

Much of our understanding of IF function has relied on comparisons between wild-type and IF knockout cells.^48^ Although valuable, knockout approaches have important limitations. First, they cannot capture the acute effects of IF loss, preventing direct observation of its immediate mechanical and structural consequences. Second, knockout cells may develop compensatory adaptations that obscure the primary roles of IFs and complicate functional interpretation. Here, we leverage FilaBuster to acutely disassemble vimentin IFs in living cells, enabling us to directly track cellular phenotypes before, during, and immediately after filament disruption. Using direct IF disassembly with FilaBuster, we uncover several immediate changes in cellular phenotype.

Acute vimentin IF disassembly triggered a retraction of asymmetrical cellular extensions and a gradual cell rounding (Fig. 5a). Cells transitioned from an elongated, irregular morphology to a more isotropic, circular shape (Fig 5b). Notably, similar cell morphology transitions have been reported in cells microinjected with high concentrations of dominant negative vimentin mimetic peptides^49^. This shift in cell architecture likely reflects changes to the organization of actin and microtubule cytoskeletons, with which VIFs make frequent interactions.

**Figure 5:**
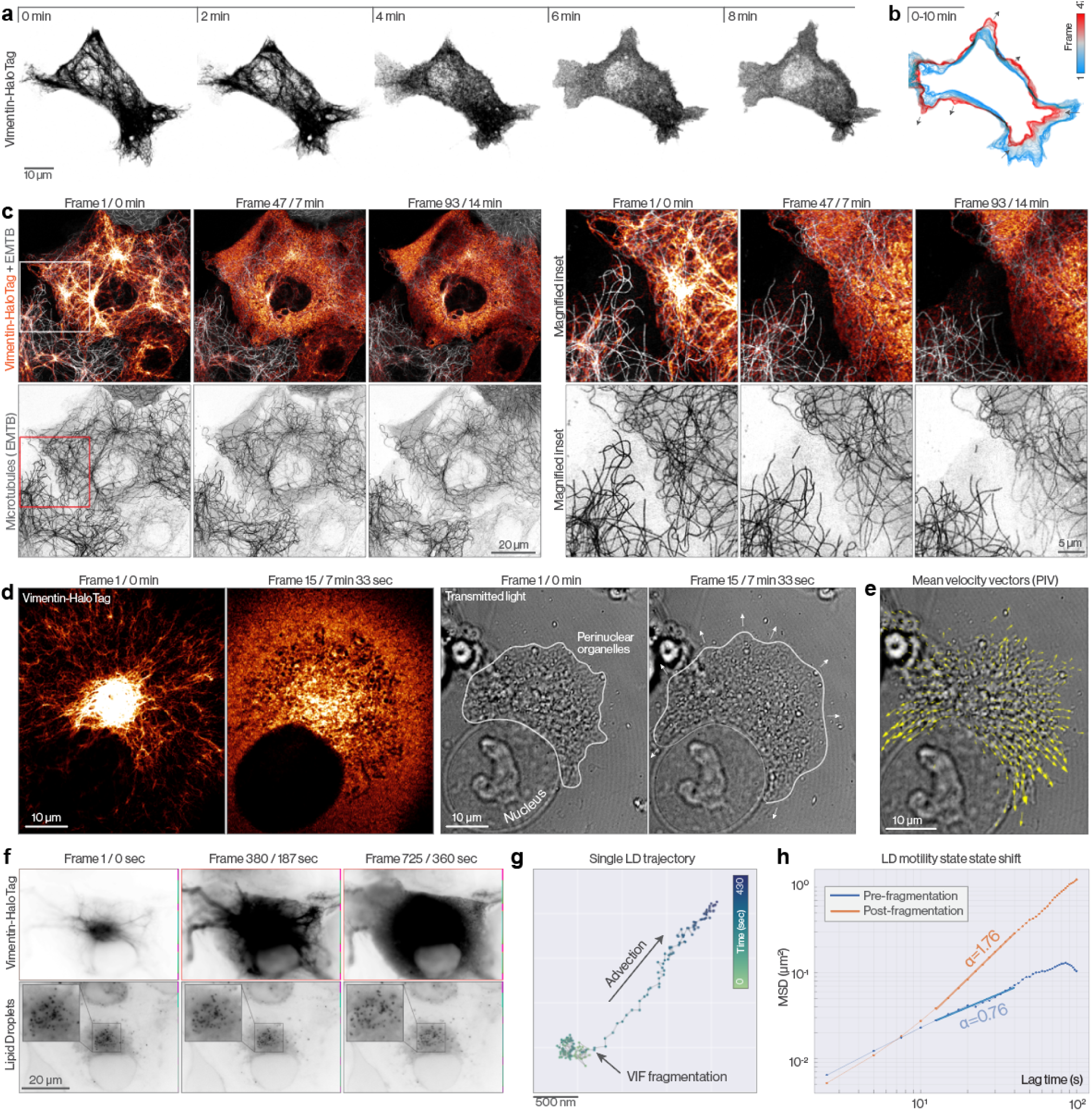
Cellular reorganization upon acute VIF fragmentation. **a.** Airyscan montage of a COS7 cell expressing Vimentin-HaloTag-JF_570_. Cellular reorganization is visible within minutes of initial VIF fragmentation. **b.** Color-coded time projection of the cell boundary from the cell indicated in a. **c.** Airyscan images of a COS-7 cell expressing vimentin-HaloTag-JF_608_ and a microtubule marker (EMTB-3×GFP). Global IF disintegration is apparent at 7 minutes, with only subtle changes to the organization and density of microtubules. **d.** Correlated fluorescence (left) and transmitted light (right) images of a COS-7 expressing vimentin-HaloTag-JF_634_ before and after VIF disintegration. White lines indicate the border of the perinuclear organellar mass. **e.** Mean velocity vectors from PIV analysis indicating motion of perinuclear organelle mass upon VIF disassembly. **f.** Widefield montage of a COS-7 expressing vimentin-HaloTag-JFX_650_ (top) and labeled with Bodipy 493/503 to visualize lipid droplets (LD, bottom). At t=3 min, the 642 nm laser power was increased, driving global VIF fragmentation and concomitant LD radial motion. **g.** Position of a representative LD over the entire movie, with arrows indicating moment of VIF disassembly. **h**. MSD of LD trajectories either before (blue) or after (red) light-mediated VIF fragmentation. Alpha values for each MSD plot are included for reference.

Indeed, VIF fragmentation elicited immediate responses in the actin cytoskeleton (Video 9). In many cells, we observed a rapid increase in actin retrograde flow, driving centripetal movement of fragmented IFs toward the perinuclear region. In other cells, actin dynamics remained largely unchanged. Over longer timescales post-fragmentation, we observed VIF fragments accumulating along stress fibers and focal adhesions. Notably, vimentin droplets that form upon recovery from hypotonic stress or expression of the vimentin Y117L ULF mutant display similar stress fiber localization.^50^ Stress fibers that became heavily decorated with VIF fragments often destabilized at later time points, resulting in rapid cell shape changes (Video 9), an effect we attribute to either loss of structural scaffolding provided by intact IF networks or from direct interactions between VIF fragments and actin filaments, which could displace or inhibit actin-binding and regulatory proteins.

We next examined the immediate effects of acute vimentin IF disassembly on microtubules, motivated by extensive literature describing their close physical and functional interactions.^51–55^ VIFs globally align with microtubules and rely on microtubule motor-based transport for proper positioning within the cytoplasm. The extent to which microtubule networks rely on VIFs for structural support is less clear. Indeed, microtubule networks appear structurally normal in vimentin knockout cells,^51^ leading to the prevailing view that microtubules do not require IFs for stability. However, vimentin 1A mimetic peptide microinjection was observed to induce dramatic microtubule depolymerization.^49^ Here, taking advantage of the spatiotemporal control afforded by the FilaBuster approach, we discover a nuanced relationship that reconciles these seemingly contrasting observations. In cells with sparse vimentin networks, IF disruption had little to no effect on microtubule integrity (Fig 5c). By contrast, in cells with dense, highly entangled vimentin IF networks, acute IF disassembly triggered rapid microtubule destabilization and fragmentation (Video 10).

These findings indicate a conditional dependence of microtubules on vimentin IFs: although microtubule networks can form normally in the chronic absence of vimentin (as observed in knockout cells), their stability becomes critically reliant on IFs when the two filament systems are extensively co-aligned and interconnected. This conditional dependence highlights a previously unappreciated functional coupling between these cytoskeletal systems—one that is uniquely revealed by FilaBuster-mediated acute perturbation but easily overlooked using traditional knockout approaches.

### Acute Vimentin IF disassembly triggers rapid redistribution of perinuclear organelles

Vimentin IFs form dense perinuclear networks that constrain organelles and regulate their positioning. While previous studies have shown that chronic depletion of vimentin leads to changes in average organelle distributions across populations,^56,57^ the direct effect of rapid and selective VIF disassembly on organelle position and dynamics has never been observed and analyzed in real time.

Here, using vimentin-HaloTag-JF_634_, we irradiated a perinuclear VIF density with 639 nm illumination and examined the effect on organelles by gentle transmitted light imaging. Upon threshold illumination levels, VIF fragments rapidly surged outwards in a radially propagating wave, traveling at ∼500 nm/sec (Fig S13a-c). Organelles that were previously entangled within the dense VIF network simultaneously lurched outwards, undergoing radial advection, resulting in more uniform membrane mass distribution within the cytoplasm and confirming a direct mechanical role for VIFs in subcellular organelle positioning (Fig 5d-e). Single-particle tracking of perinuclear, VIF-embedded lipid droplets further elucidated this effect, wherein we observed a dramatic motility state transition from subdiffusive (α=0.76) to directed outward advection (α=1.76) immediately following VIF disruption (Fig. 5f-h, Fig. S13d-e). These observations demonstrate that vimentin IFs function as a mechanical barrier, confining organelles within the perinuclear region. Acute loss of this barrier induces a rapid, advective redistribution of intracellular components, an effect masked in traditional knockout models where IF loss occurs gradually over time.

### Controlled disassembly of GFAP, desmin, peripherin, and cytokeratin filaments by activation of a HaloTag-targeted rhodamine dye

Given the shared susceptibility to oxidant-mediated disassembly reported across IF subtypes,^13,24,58^ we initially hypothesized that FilaBuster could be a universal approach for IF fragmentation, wherein filament specificity would simply be defined by the identity of the HaloTag targeting protein (i.e. IF protein or nanobody). Having thus far established this approach using VIFs as a simple proof-of-concept model system, we next evaluated whether FilaBuster could in fact be generalized to other IF subtypes.

To this end, we expressed C-terminal HaloTag fusions of additional Type III IF proteins: GFAP, desmin, and peripherin. We then electroporated these constructs into either COS-7 or U2-OS cells, labeled with optimal rhodamine HTLs based on our earlier dye benchmarking studies (Fig 2e), and acquired Airyscan images confirming that all fusions assembled phenotypically normal networks. Next, upon increased 561 nm illumination, we observed robust IF fragmentation in all cases, with similar kinetics and fragmentation patterns to what we had observed with our proof-of-concept experiments with VIFs (Fig. 6a-c, Video 11).

**Figure 6:**
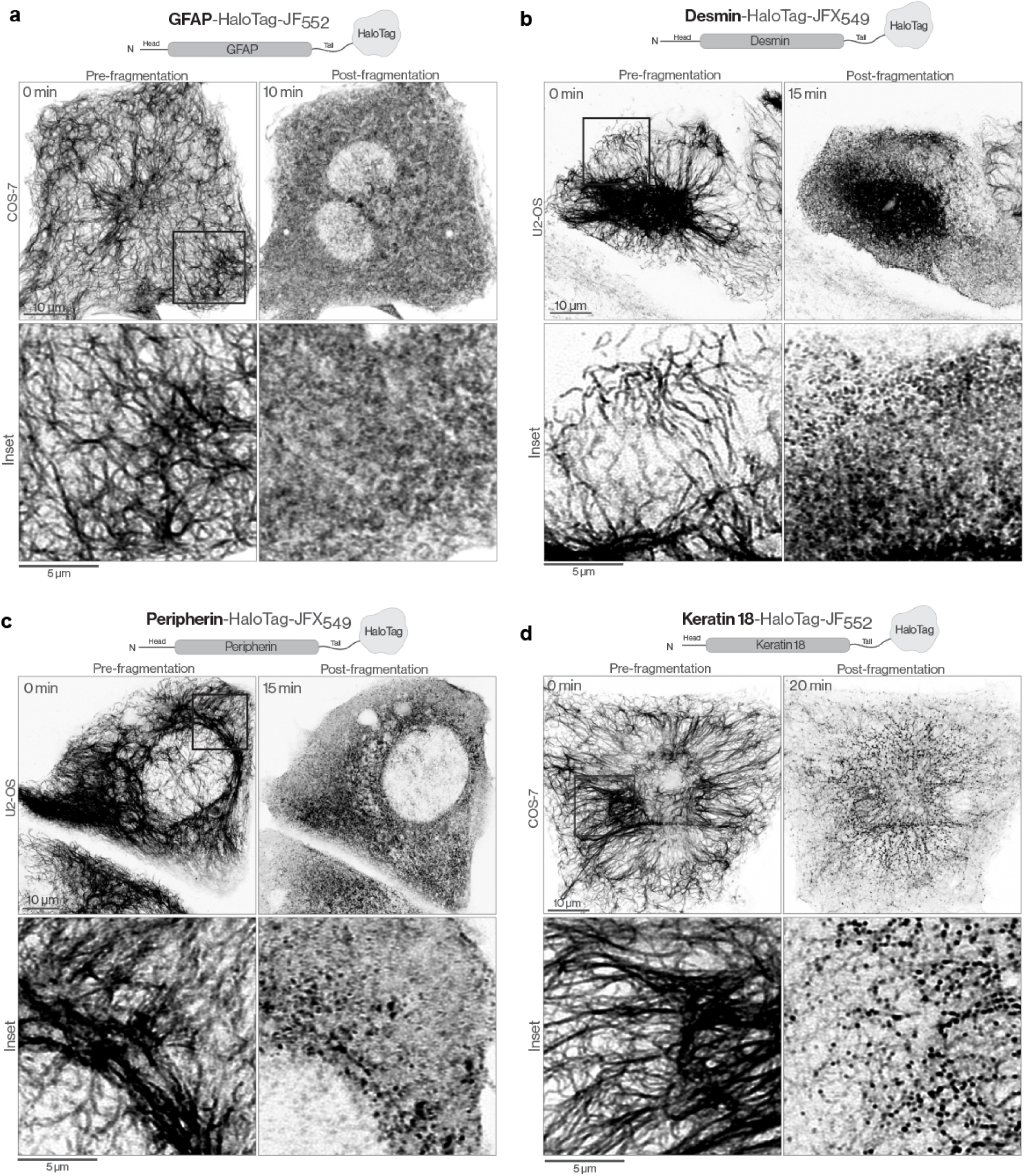
FilaBuster is a generalizable approach for IF fragmentation. **a.** Airyscan images of a COS-7 expressing GFAP-HaloTag-JF_552_ before and after 561 nm light-induced GFAP filament disintegration. **b-c.** Airyscan images of U2-OS cells expressing Desmin-HaloTag-JFX_549_ (b) or Peripherin-HaloTag-JFX_549_ (c) before and after light-induced filament disintegration. **d.** Airyscan images of a COS-7 expressing Keratin18-HaloTag-JF_552_ before and after light-mediated disassembly.

As Type III IF proteins frequently form mixed filament networks, we next tested whether targeting one type III IF protein with FilaBuster would also induce fragmentation of a separately labeled, co-expressed type III. In cells co-expressing vimentin-HaloTag-JF_552_ and Desmin-EGFP, irradiation of the HaloTag ligand triggered disassembly of both networks (Video 11). Similarly, focal 639 nm irradiation of GFAP-HaloTag-JFX_650_ induced simultaneous fragmentation of vimentin-mApple labeled VIFs (Video 11). Notably, in both cases, directly irradiating the conventionally-tagged IF alone had no effect on filament stability, confirming that disassembly only occurs when a photosensitizer ligand is activated.

Finally, we examined whether FilaBuster could be extended to other cytoplasmic IF classes. As proof of principle, we focused on the abundant and well characterized type I cytokeratin, keratin 18. We electroporated COS-7 cells with a keratin 18-HaloTag construct, labeled with JF_552_-HTL, and observed a dense cytoplasmic network, consistent with efficient heterodimerization with endogenous type II keratins. Upon sustained 561 nm irradiation, we observed clear loss of keratin IFs; however, the disassembly process appeared qualitatively distinct from that of type III IFs. Rather than abruptly disintegrating into minimal fragments scattered throughout the cytoplasm, as seen with vimentin (Fig. 4a-c), keratin filaments gradually developed discrete, granular structures along their length, reminiscent of the structural reorganization reported upon orthovanadate treatment^59^. Over several minutes, these puncta increased in size and intensity, ultimately depleting keratin from the filament network until only these structures remained (Fig. 6d, Video 11).

## Discussion

In this study, we introduced FilaBuster, a generalizable strategy for rapid, specific, and spatiotemporally controlled IF disassembly. The approach consists of three main steps: (1) targeting a HaloTag fusion protein to native IFs, (2) labeling with a cell-permeable photosensitizer HaloTag ligand, and (3) activating the photosensitizer with light. We show that this simple, modular workflow enables irreversible photochemical IF destabilization with remarkable speed, specificity, and reliability.

To validate this approach, we first applied FilaBuster to vimentin IFs as a model system. As a means of targeting HaloTag to VIFs (step 1), we designed a vimentin-HaloTag fusion protein that coassembles with endogenous filaments. After confirming proper filament incorporation with fluorescence microscopy and cryo-ET, we labeled cells with the photosensitizer dye JF_570_-HTL (step 2) and briefly irradiated with low-power 561 nm light (step 3), driving rapid and complete disassembly of the cellular VIF network. Having established proof-of-concept, we next explored the versatility of FilaBuster.

First, we introduced and validated a novel red-shifted JaneliaFluor photosensitizer dye, JF_634_, enabling acute VIF fragmentation with far red laser lines. Next, we validated a vimentin nanobody-HaloTag fusion as an alternative labeling strategy capable of binding and photo-fragmenting native filaments without the need for vimentin subunit tagging or ectopic vimentin expression. From there, we screened a panel of HaloTag ligands, confirming that conventional rhodamine dyes—despite not being designed as photosensitizers—can, under sufficient photon flux, drive photochemical IF disassembly, allowing both imaging under low power and filament disassembly using the same label. Finally, we benchmarked different illumination conditions, determining light dose thresholds required for efficient IF disassembly for different HTLs, and comparing parameters for focal versus global VIF disintegration.

Using these approaches, we examined the immediate consequences of acute vimentin IF disassembly, identifying rapid changes in cell shape, cytoskeletal organization, and organelle positioning. Extensive controls confirmed that FilaBuster is specific, reproducible, and non-toxic under conditions that effectively dismantle VIFs, with cells remaining viable 20 hours post-fragmentation (Supplementary Fig. 12c). We further demonstrated that FilaBuster extends beyond vimentin, enabling selective disruption of GFAP, desmin, peripherin, and keratin-18 networks. These findings establish FilaBuster as a powerful tool for disrupting IF networks in space and time, filling a long-standing methodological gap for achieving acute, selective, and tunable IF perturbation in living cells.

To understand why FilaBuster is so effective—and why IFs, among all cytoskeletal components, are particularly susceptible to this strategy—we turned to the natural roles that IFs play in regulating redox homeostasis. Beyond their established functions in mechanical support and structural organization, cytoplasmic IFs are increasingly recognized as passive buffers against oxidative damage.^60,61^ Their extensive, filamentous network within the cytoplasm, coupled with their remarkable capacity to undergo a wide range of oxidative and electrophilic modifications^62^, positions them as a frontline sink for potentially toxic ROS. Moreover, their modular organization, ongoing subunit exchange,^63^ and continuous self-repair ^64^, allows gradual turnover of modified subunits during mild oxidative stress without complete filament destabilization. However, when the oxidative load exceeds a critical threshold, IFs disassemble abruptly, sacrificing themselves to protect more vulnerable cellular structures.^65^ Indeed, mitochondria located in regions rich in vimentin filaments have been shown to exhibit greater resilience to oxidative stress, ^66^ lending further support to this protective buffering role.

FilaBuster transforms this general oxidative vulnerability into a highly specific and optically controlled mechanism for filament disassembly. Rather than flooding the cytoplasm with reactive oxidants, it embeds a light-triggered disassembly mechanism directly into the filament network. Cryo-electron tomography revealed that HaloTags are tethered just nanometers from the filament surface— within the effective damage radius of short-lived ROS such as singlet oxygen—and regularly positioned along the filament to ensure maximal coverage. Because the HaloTag fusion co-assembles with wild-type IF subunits, each photosensitizer can affect neighboring untagged subunits, further amplifying the effect. Under normal conditions, the network behaves indistinguishably from its endogenous counterpart—but upon activation, it disassembles with exceptional efficiency.

We believe this is the first strategy that enables rapid, selective, and optically controlled disassembly of IFs in live cells. Its versatility, modularity, and compatibility with a wide spectrum of ligands offer a path forward for dissecting the compartmentalized roles of IFs in processes ranging from signaling and trafficking to stress adaptation and mechanical resilience. By finally equipping the field with a tool capable of acute and localized IF perturbation, FilaBuster opens the door to a deeper, more spatially resolved understanding of one of the cell’s most enigmatic cytoskeletal systems.

## Methods

### Plasmids

Lentiviral vectors for vimentin-HaloTag, Vimentin-C328A-HaloTag, vimentin-C328H-HaloTag, vimentin-SnapTag, vimentin-mNeonGreen, vimentin-SuperNova2, vimentin-mStayGold, desmin-HaloTag, and peripherin-HaloTag were cloned by VectorBuilder. In all cases, the tag was added to the c terminus of human vimentin, desmin, or peripherin via a six residue (GSGSGS) linker. GFAP-HaloTag (Addgene, 169474) and Vimentin-mEmerald (Addgene, 54299) were obtained from Addgene. The vimentin-chromobody (Vimentin-VHH-TagGFP2) was purchased from Chromotek (now ProteinTech, vcg). Vimentin-VHH-HaloTag was generated by synthesizing coHaloTag - a cysteine-free variant of HaloTag7 with C61T and C262V amino acid substitutions, and subcloning into the Vimentin-VHH-TagGFP2 plasmid using HindIII and NotI restriction sites (genscript). As a result, coHaloTag is linked to the c-terminus of the vimentin VHH/nanobody via a 22 amino acid linker. Keratin18-HaloTag was obtained from the (Janelia Cell and Molecular Biology core facility).

### Cell culture

COS-7 (ATCC), PTK2 (ATCC), RPE-1 (ATCC), MEF (WT and Vim-/-), HEK293T, U2-OS (ATCC), Vimentin-HaloTag knock-in U2-OS (courtesy of Stefan Jakobs), and NIH3T3 (ATCC) were maintained in DMEM (Corning) with 4.5 g/L glucose supplemented with 10% fetal bovine serum, 2 mM Glutamax, and 100 U/mL penicillin and 100 µg/mL streptomycin. Normal human dermal fibroblast (NHDF MatTek; NHDF-CRY-NEO) were cultured in NHDF growth medium (MatTek).

All cells were maintained in a 37C, 5% CO_2_ incubator. For imaging experiments, cells were seeded on either 35 mm #1.5 glass-bottom dishes (MatTek, P35G-1.5-C-C) or 12 well #1.5 glass-bottom plates (CellViz; P12-1.5H-N) coated with either fibronection or matrigel (500 µg/ml). Transient transfection was performed using either FuGene HD & FuGene 4k (Roche). Nucleofection was performed using an Amaxa Nucleofector and Solution SE for COS-7 and U2-OS. Imaging was performed in complete culture medium and cells were maintained at 37 ° / 5% CO_2_ throughout all live imaging experiments. Drug treatments were formed using 10 µM nocodazole diluted in DMSO (Millipore Sigma, M1404), 10 µM taxol diluted in DMSO (Millipore Sigma, PHL89806), or 3 µM Latrunculin A diluted in DMSO. (Millipore Sigma, L5163). Cells were incubated with HaloTag ligands at 250 nM for 30-60 minutes and washed 3x in complete media before imaging. Cells were incubated with CellROX Deep Red (Invitrogen, C10422) at 5 µM concurrently with HTL labeling.

### Stable cell lines

Vimentin-mEmerald knock in COS-7: mEmerald was knocked into the C terminal of vimentin in COS-7 cells using CRISPR-Cas9. Guide RNA (gRNA) was designed using CHOPCHOP and screened for off target cutting and efficiency using CCTop. Both gRNA and Alt-R™ S.p. HiFi Cas9 were procured from IDT, while a ssDNA donor and oligo containing a tCTS sequence were synthesized by Genscript. The ribonucleoprotein (RNP) complex was formed with 150pmol sgRNA and 125pmol of IDT’s Alt-R S.p.Cas9-GFP V3 protein. The RNP was incubated at room temperature for 20 minutes. The GenExact ssDNA donor template was annealed to a DNA oligo to form partial dsDNA structure by combining ssDNA with oligo at a 1:4 molar ratio and incubating at 70 °C for 5 minutes and gradually decreasing the temperature to 4 °C at a rate of 5°C per 5 minutes. 1 µg (3.37pmol) of annealed template was used in the nucleofection along with 120 pmol of IDT Cas9 Electroporation Enhancer.The complexed RNP was combined with the annealed donor according to manufacturers’ protocols. COS-7 cells were nucleofected using the Lonza 4D-Nucleofector platform and recovered in media containing IDT’s HDR enhancer for 24 hours before isolating single cell clones via FACS sorting. Clones were screened with PCR and nanopore sequencing to confirm the presence of a correct heterozygous insertion and intact wild-type allele.

Vimentin-HaloTag COS-7, RPE-1, HEK293T, NIH-3T3, MEF, and NHDF stable lines were generated by lentiviral transduction. In brief, parent cells were seeded in 6 well dishes and incubated with different doses (0.5 - 2 µL) of high titer (>10^8^ TU/ml) lentivirus for 24 hours. Cells were then washed twice in fresh media, and expression levels/cell viability was monitored over several days before selecting polyclonal lines with intermediate expression levels.

### Immunocytochemistry

Polyclonal chicken anti-Vimentin (Encor biotech, CPCA-Vim) and Goat anti-chicken IgY Alexa 405-plus (ThermoFisher, A48260) were used for staining vimentin (Fig S1a). Fixation was performed with 4% paraformaldehyde in PBS (electron microscopy sciences), followed by 3× washes in PBS, permeabilization in 0.25% Triton X-100 for 1h, and incubation overnight with primary antibodies in 0.1% TritonX100 in PBS. Samples were washed 3× in PBS, and incubated for one hour with secondary antibodies in 0.1% Triton X-100 before imaging in PBS.

### Dye labeling and HTL screen

Stable vimentin-HaloTag MEFs were seeded in 12-well glass-bottom plates (CellViz). At ∼75% confluency, cells were incubated with HaloTag ligand (250 nM, 0.5-1h) in complete medium. Cells were then washed 3x in complete DMEM and transferred to a temperature and CO2 controlled stage for widefield epifluoresence microscopy. Time-lapse movies were acquired at one frame per second with a 63×/1.4NA objective lens and 1.6 optovar. At each frame, the sample was irradiated for 100 ms using either 488 nm, 561 nm, or 640 nm laser lines (see power measurements and irradiance/energy calculations below). Timelapse movies were acquired for 600 frames at 1 Hz, but were terminated earlier if all cells in the field of view had undergone IF fragmentation. Movies were then blinded and manually analyzed to determine the latency (number of seconds / irradiation cycles) to initial VIF fragmentation.

### Imaging systems

#### Airyscan confocal microscopy

Airyscan microscopy was performed on either a Zeiss 880 or Zeiss 980 using an Axio Observer Z1 inverted microscope. Samples were illuminated with 488 nm/561 nm/633 nm laser lines on the Zeiss880 and 488 nm / 561 nm / 639 nm laser lines on the Zeiss 980. In both cases, samples were imaged with a Plan-Apochromat 63x/1.4 Oil DIC M27 objective lens. Samples imaged on the 980 were acquired using Airyscan 2.0 SR mode and automatically processed in Zen Blue (Zeiss).

#### Confocal laser power and irradiance measurements

Laser power at the sample plane was measured using a Newport 1936-R power meter coupled with a calibrated silicon photodiode sensor (Newport 818 series). Because the microscope configuration was inverted, the sensor was placed directly at the sample position above the objective lens. To calculate irradiance, we determined the effective beam area by performing a spot bleach experiment using the 561 nm laser line at 100% power on fixed Alexa Fluor 555-immunostained samples. Images were inverted, and a Gaussian function was fit to the intensity profile obtained from a one-pixel-wide line scan across the bleached region. The full width at half maximum (FWHM) calculated from the fit FWHM = 2√(2 ln 2)·σ) was 488.1 nm, corresponding to a beam radius of approximately 244.05 nm, consistent with the theoretical diffraction-limited spot size.

Laser power measurements were taken at various laser power percentage settings in bidirectional scanning mode, the mode used during imaging experiments. Due to AOTF-mediated laser blanking during scanner retrace periods, these measurements represent time-averaged power, slightly underestimating continuous laser output. To quantify this difference, we additionally measured power using “spot mode,” with the laser continuously unblanked and focused at the sample plane for five seconds. Bidirectional scanning mode measurements were found to be 85.5% ± 1.0% (mean ± SD) of those obtained in spot mode. For example, at 1% laser power, bidirectional scanning yielded 3.16 µW, while spot mode yielded 3.72 µW.

Measured laser power values were used to generate a calibration curve through least-squares linear regression, focusing on the 0.01%–1% power range, which encompassed the settings most frequently used during imaging experiments. This calibration allowed interpolation of laser power across all experimental settings. Irradiance (W/cm²) was calculated by dividing measured or interpolated power by the effective beam area determined from the bleaching experiments. The energy dose per pixel (J/cm²) was subsequently determined by multiplying irradiance by the pixel dwell time. Similar procedures were performed to measure laser power

#### Widefield microscopy

Widefield epifluorescence, total internal reflection fluorescence (TIRF) microscopy, and Lattice SIM were performed on a Zeiss Elyra 7 using an AxioObserver inverted microscope. Samples were illuminated with 500 mW 640 nm, 561 nm, and 488 nm laser lines using a Plan-Apochromat 63x/1.4 Oil DIC M27 objective lens and 1.6X optovar. Emission was collected with dual EM-CCD cameras.

#### Widefield Laser Power and Irradiance Measurements

Widefield laser power measurements were performed using a Thorlabs PM100D power meter with an S170C microscope slide-format photodiode sensor. All measurements were performed in Laser WF mode using the same imaging configuration as in experiments, including the 63× /1.4 NA objective lens and 1.6× optovar. To determine the effective beam area, we first performed a photobleaching experiment using the 561 nm laser at 100% power on a fixed sample immunostained with Alexa Fluor 555. A 3×3 stitched image was acquired after bleaching to visualize the full extent of the bleached region. The bleached area was segmented and quantified from the stitched image to calculate the projected beam area of 13273 µm² at the sample plane.

Laser power was measured at 0.2%, 2%, and 20% settings for each of the three 500 mW laser lines (488 nm, 561 nm, and 642 nm). For the 488 nm line, measured power values were 0.58 mW (0.2%), 2.9 mW (2%), and 20 mW (20%), corresponding to irradiances of 4.4 W/cm², 21.8 W/cm², and 150.7 W/cm², respectively. For the 561 nm line, measured power values were 0.5 mW (0.2%), 2.5 mW (2%), and 29 mW (20%), corresponding to irradiances of 3.8 W/cm², 18.8 W/cm², and 218.5 W/cm², respectively. For the 642 nm line, measured power values were 0.5 mW (0.2%), 2.7 mW (2%), and 20 mW (20%), corresponding to irradiances of 3.8 W/cm², 20.3 W/cm², and 150.7 W/cm², respectively.

#### Cryo-ET of detergent-treated MEFs

MEFs expressing Vimentin-Halotag were cultured in DMEM (Sigma-Aldrich, D5671), supplemented with 10% FBS (Sigma-Aldrich, F7524), 2 mM l-glutamine (Sigma-Aldrich, G7513) and 100 μg ml–1 penicillin–streptomycin (Sigma-Aldrich, P0781), at 37 °C and 5% CO_2_ in a humidified incubator.

MEFs expressing vimentin-HaloTag were cultured to ∼80% confluency on glow-discharged holey carbon EM grids (R2/2, Au 200 mesh; Quantifoil). MEF grids were washed in 37°C PBS, 2 mM MgCl_2_ for 5 s before being transferred for 30 s into 37°C PBS, 0.1% Triton X-100, 10 mM MgCl_2_, 600 mM KCl, cOmplete protease inhibitors. Grids were washed in PBS, 2 mM MgCl_2_ for 10 s and then incubated for 30 minutes in PBS, 2 mM MgCl_2_, 2.5 units/µl benzonase, 40 ng/µl DNase I at room temperature. After the enzymatic treatment, the grids were washed for 5 s in PBS, 2 mM MgCl_2_ and 3 µl of 10 nm fiducial gold markers (Aurion) was applied. The grids were immediately blotted for 5 s from the reverse side and plunge frozen in liquid ethane using a manual plunge freezer.

Tilt-series were collected using a 300 keV Titan Krios with a K2 Summit detector and Quantum energy filter set to a slit width of 20 eV. Data acquisition was performed using SerialEM^67^ and PACE-tomo ^68^ in low-dose mode at ×81.000 magnification, pixel size 1.72 Å, over a ± 60° dose symmetric tilt range with 3° increments, −4 μm defocus, and a cumulative dosage of ∼150 e–/Å2. Images were drift-corrected using MotionCor2^69^ (Zheng et al. 2017) and aligned using fiducial markers in IMOD^70,71^. Tomograms were reconstructed and binned 6 times (pixel size 10.8 Å) with weighted back-projections and a SIRT-like filter was used for enhancing image contrast. HaloTags were observed as spherical densities with a 3-4 nm diameter, fitting the expected size, and distributed within a 30 nm distance around well-assembled vimentin filaments (diameter ∼11 nm). A subset of vimentin filaments and HaloTags were manually picked with Napari^72^ and used to train convolutional neural network models in crYOLO^73,74^. Vimentin filaments and halotags were predicted using neural network models and manually curated in five tomograms. MATLAB scripts were used to calculate vimentin filament trajectories through tomogram volumes by linear interpolation of vimentin coordinates at filament centers. Vimentin interpolations were used to calculate the distance and distributions of HaloTags to and along vimentin filaments. Tomogram images and vimentin-halotag 3D model were prepared in IMOD^70,71^ and Blender 3D.

#### Dye synthesis

Commercial reagents were obtained from reputable suppliers and used as received. All solvents were purchased in septum-sealed bottles stored under an inert atmosphere. All reactions were sealed with septa through which a nitrogen atmosphere was introduced unless otherwise noted. Reactions were conducted in round-bottomed flasks or septum-capped crimp-top vials containing Teflon-coated magnetic stir bars. Heating of reactions was accomplished with a silicon oil bath or an aluminum reaction block on top of a stirring hotplate equipped with an electronic contact thermometer to maintain the indicated temperatures.

Reactions were monitored by thin layer chromatography (TLC) on precoated TLC glass plates (silica gel 60 F_254_, 250 µm thickness) or by LC/MS (Phenomenex Kinetex 2.1 mm × 30 mm 2.6 μm C18 column; 5 μL injection; 5–98% MeCN/H_2_O, linear gradient, with constant 0.1% v/v HCO_2_H additive; 6 min run; 0.5 mL/min flow; ESI; positive ion mode). TLC chromatograms were visualized by UV illumination or developed with *p*-anisaldehyde, ceric ammonium molybdate, or KMnO_4_ stain. Reaction products were purified by flash chromatography on an automated purification system using pre-packed silica gel columns or by preparative HPLC (Phenomenex Gemini NX-C18 30 × 150 mm 5 μm column). Analytical HPLC analysis was performed with a Phenomenex Gemini NX-C18 4.6 × 150 mm 5 μm column under the indicated conditions.

NMR spectra were recorded on a 400 MHz spectrometer. ^1^H and ^13^C chemical shifts were referenced to TMS or residual solvent peaks. Data for ^1^H NMR spectra are reported as follows: chemical shift (δ ppm), multiplicity (s = singlet, d = doublet, t = triplet, q = quartet, dd = doublet of doublets, m = multiplet), coupling constant (Hz), integration. Data for ^13^C NMR spectra are reported by chemical shift (δ ppm) with hydrogen multiplicity (C, CH, CH_2_, CH_3_) information obtained from DEPT spectra.

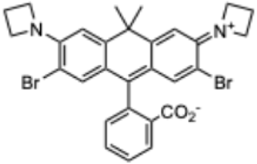

#### JF_634_ (2′,7′-dibromo-JF_608_)

JF_608_ (73 mg, 0.167 mmol) was taken up in CH_2_Cl_2_ (2 mL) and cooled to 0 °C. *N*-Bromosuccinimide (59.6 mg, 0.335 mmol, 2 eq) in DMF (1 mL) was added dropwise over 5 min, and the reaction was stirred at 0 °C for 15 min. It was subsequently diluted with saturated NaHCO_3_ and extracted with EtOAc (2×). The combined organic extracts were dried over anhydrous MgSO_4_, filtered, and concentrated *in vacuo*. Silica gel chromatography (0–50% EtOAc/toluene, linear gradient) afforded 79.3 mg (80%) of the title compound as a pale blue solid. ^1^H NMR (CDCl_3_, 400 MHz) δ 8.04 – 8.00 (m, 1H), 7.65 (td, *J* = 7.4, 1.4 Hz, 1H), 7.60 (td, *J* = 7.4, 1.2 Hz, 1H), 7.09 – 7.05 (m, 1H), 6.67 (s, 2H), 6.63 (s, 2H), 4.18 – 4.07 (m, 8H), 2.29 (p, *J* = 7.3 Hz, 4H), 1.80 (s, 3H), 1.71 (s, 3H); ^13^C NMR (CDCl_3_, 101 MHz) δ 170.3 (C), 154.1 (C), 149.8 (C), 145.5 (C), 135.1 (CH), 133.4 (CH), 129.7 (CH), 126.9 (C), 125.5 (CH), 123.9 (CH), 123.6 (C), 111.6 (CH), 106.9 (C), 86.2 (C), 54.4 (CH_2_), 38.4 (C), 35.6 (CH_3_), 32.2 (CH_3_), 16.8 (CH_2_); Analytical HPLC: t_R_ = 17.2 min, >99% purity (10–95% MeCN/H_2_O, linear gradient, with constant 0.1% v/v TFA additive; 20 min run; 1 mL/min flow; ESI; positive ion mode; detection at 650 nm); MS (ESI) calcd for C_29_H_27_Br_2_N_2_O_2_ [M+H]^+^ 595.0414, found 595.2.

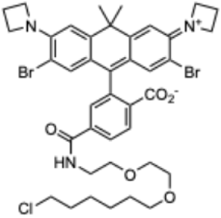

#### JF_634_–HaloTag ligand (2′,7′-dibromo-JF_608_–HaloTag ligand)

JF_608_–HaloTag ligand (20 mg, 0.029 mmol) was taken up in CH_2_Cl_2_ (1 mL) and cooled to 0 °C. *N*-Bromosuccinimide (10.4 mg, 0.058 mmol, 2 eq) in DMF (0.5 mL) was added dropwise over 5 min, and the reaction was stirred at 0 °C for 15 min. It was subsequently diluted with saturated NaHCO_3_ and extracted with EtOAc (2×). The combined organic extracts were washed with brine, dried over anhydrous MgSO_4_, filtered, and concentrated *in vacuo*. Purification by reverse phase HPLC (50–95% MeCN/H_2_O, linear gradient, with constant 0.1% v/v TFA additive) afforded 10.5 mg (43%) of the title compound as a pale blue solid. ^1^H NMR (CDCl_3_, 400 MHz) δ 7.99 (dd, *J* = 8.0, 0.7 Hz, 1H), 7.89 (dd, *J* = 8.0, 1.4 Hz, 1H), 7.39 (dd, *J* = 1.4, 0.8 Hz, 1H), 6.77 (t, *J* = 4.6 Hz, 1H), 6.58 (s, 2H), 6.54 (s, 2H), 4.06 (t, *J* = 7.3 Hz, 8H), 3.60 – 3.52 (m, 6H), 3.50 – 3.47 (m, 2H), 3.45 (t, *J* = 6.7 Hz, 2H), 3.33 (t, *J* = 6.6 Hz, 2H), 2.23 (p, *J* = 7.3 Hz, 4H), 1.74 (s, 3H), 1.71 – 1.65 (m, 2H), 1.63 (s, 3H), 1.47 – 1.42 (m, 2H), 1.39 – 1.31 (m, 2H), 1.29 – 1.23 (m, 2H); Analytical HPLC: t_R_ = 15.3 min, >99% purity (30–95% MeCN/H_2_O, linear gradient, with constant 0.1% v/v TFA additive; 20 min run; 1 mL/min flow; ESI; positive ion mode; detection at 650 nm); MS (ESI) calcd for C_40_H_47_Br_2_ClN_3_O_5_ [M+H]^+^ 844.1546, found 844.3.

### Mass Spectrometry

Solvents and Chemicals: Water (Optima, W6–4), acetonitrile (ACN, Optima, A9554), methanol (Optima, A454SK-4), formic acid (Pierce, PI28905), Acetone (A949-1, Fisher) RapiGest SF Surfactant (Waters,186001861). Dithiothreitol (DTT, R0861, Thermo scientific), Trifluoroacetic acid (TFA, 302031–100ML), Iodoacetamide (IAA, 407710) and ammonium bicarbonate (40867)) were purchased from Sigma Aldrich. Trypsin (V5113) was purchased from Promega.

Protein Digestion and Desalting: The concentration of the proteins from each cell lysate was assessed through BCA assay. The same amount of proteins from each sample were then precipitated with cold acetone at 1:4 (v:v) ratio, -20 °C overnight. Protein pellets were centrifuged and washed with cold acetone the next day and the residue acetone was air dried. The precipitated protein pellets were re-dissolved using 0.5% RapiGest according to the manufacturer’s protocol. Afterwards, proteins were reduced with 5mM DTT (60°C, 30 min) and alkylated using 11mM IAA at ambient temperature in the dark for 30 minutes. Trypsin was added at a protein-enzyme ratio of 50:1 for overnight digestion at 37 °C. The digestion was quenched by adding 10% TFA to pH ∼1. The digest was further incubated at 37 °C for 45 min to degrade the RapiGest. The solutions were then centrifuged at 14 000 rpm for 10 min and the supernatant peptides were collected. Desalting of the peptides was performed using C18 ZipTip (Millipore, ZTC18M096) and the eluents were dried using SpeedVac (Thermo Scientific). The samples were stored at –80 °C before being re-suspended in 0.1% formic acid for LC-MS/MS analysis.

LC-MS/MS Analysis: LC separation was performed on a Vanquish Neo System (Thermo Scientific) with an IonOptik Aurora Ultimate C18 column (25cm length × 75 μm inner diameter × 1.7 μm particle size) at 50 °C and 300 nL/min flow rate. Mobile phase A consisted of 0.1% formic acid in water. Mobile phase B consisted of 0.1% formic acid in 80% ACN. Eluting peptides were ionized by electrospray ionization and then analyzed by an Orbitrap Ascend Tribrid mass spectrometer (tune version 4.1.4244, Thermo Scientific). Ion transfer tube temperature was set to 275 °C. Source positive ion voltage was set to 1850v. For single-shot proteomics with data dependent acquisition (DDA), the MS1 scan resolution was set to 120,000 (at m/z 200). MS1 scan range was 350–1500 m/z, AGC target was 250%, maximum injection time mode was set to auto. Precursors were isolated through quadrupole with an isolation width of 1.2 (m/z). Precursors were fragmented by HCD at an NCE of 26%. MS2 scans were acquired by ion trap using rapid rate.

Data Processing: Raw data was directly processed using Thermo Proteome Discoverer (3.0.0.757) against the homo sapiens (UP000005640) proteome fasta file with the Halo tag attached to the vimentin sequence. A maximum of one missed cleavage was allowed and cysteine carbamidomethylation (+57.0215 Da) as the fixed modification. Oxidation(M), Phosphorylation (S, T, Y) and acetylation at protein N-terminus were selected as variable modification. Percolator node was used, and the target FDR (strict) was set to 0.01. 10 ppm was used as the precursor mass tolerance and 0.2 Da for the fragment ion tolerance window. Label-free quantification information of the peptides and proteins was processed with sample runs normalized according to the total peptide amount.

### Image processing and analysis

Local image coherency, energy, and orientation were measured with the ImageJ plugin OrientationJ using cubic spline 2px gradients. To measure local coherence changes over time, coherency maps were threshold by 50% energy and mean values were plotted over time. Coefficient of Variation (CV) and normalized radial vimentin intensity were analyzed using custom MATLAB scripts. Normalized CV change computes the fractional change in CV at each frame relative to the first frame’s CV. This metric allows for easier comparison between CV changes in images with different baseline signal to noise ratios and was implemented in Python. For wide field epifluorescence movies, bleach correction with exponential fit was performed in ImageJ for dyes that displayed appreciable bleaching over the imaging window. For presentation, a subset of movies were filtered using the unsharp mask filter in Fiji using a radius of 5 px and mask weight of 0.6pt before using a three-frame sum intensity projection, increasing VIF contrast but reducing temporal resolution. GPT4 (OpenAI) was used for assistance in troubleshooting and converting pre-existing MATLAB scripts into python.

Single particle tracking (SPT) data was performed in ImageJ using the TrackMate plugin^75^. For SPT of VIF fragments, images were convolved with a Gaussian blurring filter with a kernel size of 2 px. Particle localization was performed using the Hessian detector algorithm, and the simple LAP Tracker was used to generate trajectories with the following conditions: 500 nm maximum gap closing distance, no frame gaps, and 500 nm frame-to-frame linking distance. For SPT of lipid droplets, the Laplacian of Gaussian detector with an estimated object diameter of 0.3 µm was used to identify LDs, and the simple LAP tracker was used with 200 nm linking max distance, 500 nm gap-closing max distance, and gap-closing max frame gap of 2. For VIF fragment trajectories, mean-squared displacements (MSDs) were calculated in Mathematica 14.0 and were piecewise fit to MSD(t) = ɑ for effective diffusion coefficient D, time t, and time dependence ɑ. For LDs, α values were obtained in python by fitting a linear model to the log-transformed MSD data over the 2nd through 18th lag time increments, with slope α. MSD, net displacement, and trajectories were plotted in python using the Seaborn library.

Fluorescence Recovery After Photobleaching (FRAP) measurements were performed using ImageJ by selecting ROIs around VIF fragments and quantifying mean fluorescence intensity over time using the Z-Axis Profile function. The recovery curve of mean intensity I(t) was fit using the exponential recovery function: *I*(*t*) = *A*(1 − *e*^(−*t*/τ)^) + *B* where *A* represents amplitude of fluorescence recovery, τ is characteristic recovery time, and *B* is baseline offset. Mobile fraction was calculated by normalizing amplitude *A* to the mean fluorescence intensity prior to photobleaching, and immobile fraction was determined as (1−mobile fraction).

## Acknowledgements

We thank the Janelia Cell and Molecular Biology core facility, J. Towne, M. Ramirez, and N. Rivero Ballón for technical assistance; to H. Choi, C. Gladkova, and S. Khuon for valuable discussions and critical feedback; to S. Upadhyayula, M. DeSantis, and B. Crooks for valuable discussions concerning measuring and reporting power density; to R. Patel and T. Brown, for sharing absorbance spectra of HaloTag-bound JF HTLs; and to T Stephen and S. Jakobs for sharing endogenously tagged vimentin-HaloTag U2-OS cells. We thank the Harvard Center for Biological Imaging (RRID:SCR_018673) for infrastructure and support. SBH was supported by an EMBO Postdoctoral Fellowship (ALTF 683-2023).

