## Supplementary Figures and Video Captions for "FilaBuster: A Strategy for Rapid, Specific, and Spatiotemporally Controlled Intermediate Filament Disassembly"

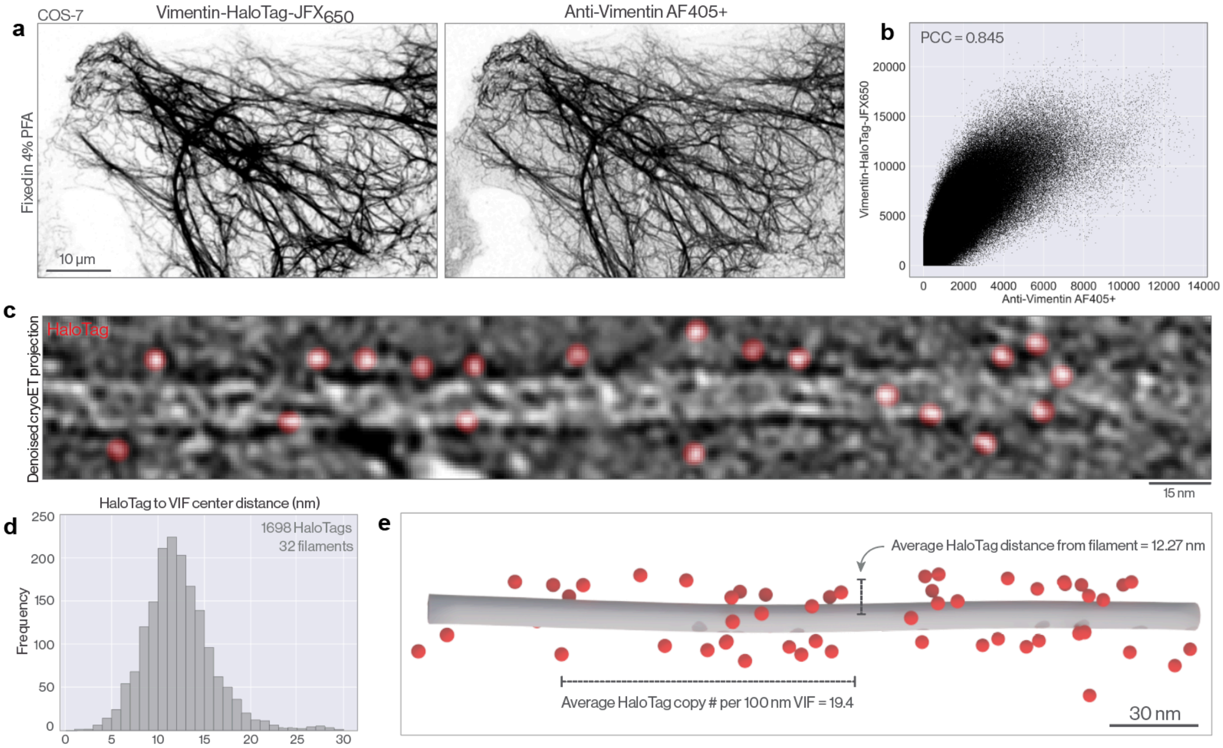


**Supplementary Figure 1: Vimentin-HaloTag incorporates into the endogenous VIF network**
**a.** Airyscan images of a chemically fixed COS-7 expressing Vimentin-HaloTag-JFX$_{650}$ (left) and immunostained for vimentin (right) **b**. Pixel-wise intensity distribution for the images shown in a with Pearson's Correlation Coefficient (PCC). **c.** Projected, denoised tomogram of a single vimentin filament in a MEF. Densities corresponding to HaloTag are pseudocolored red. **d.** Frequency distribution of HaloTag to VIF distances along 32 filaments from 5 tomograms. **e.** Three-dimensional rendering indicating position of vimentin-HaloTag incorporation into a single VIF. Rendering is annotated with mean HaloTag copy number per 100 nm and mean HaloTag distance from filament computed from the full dataset.

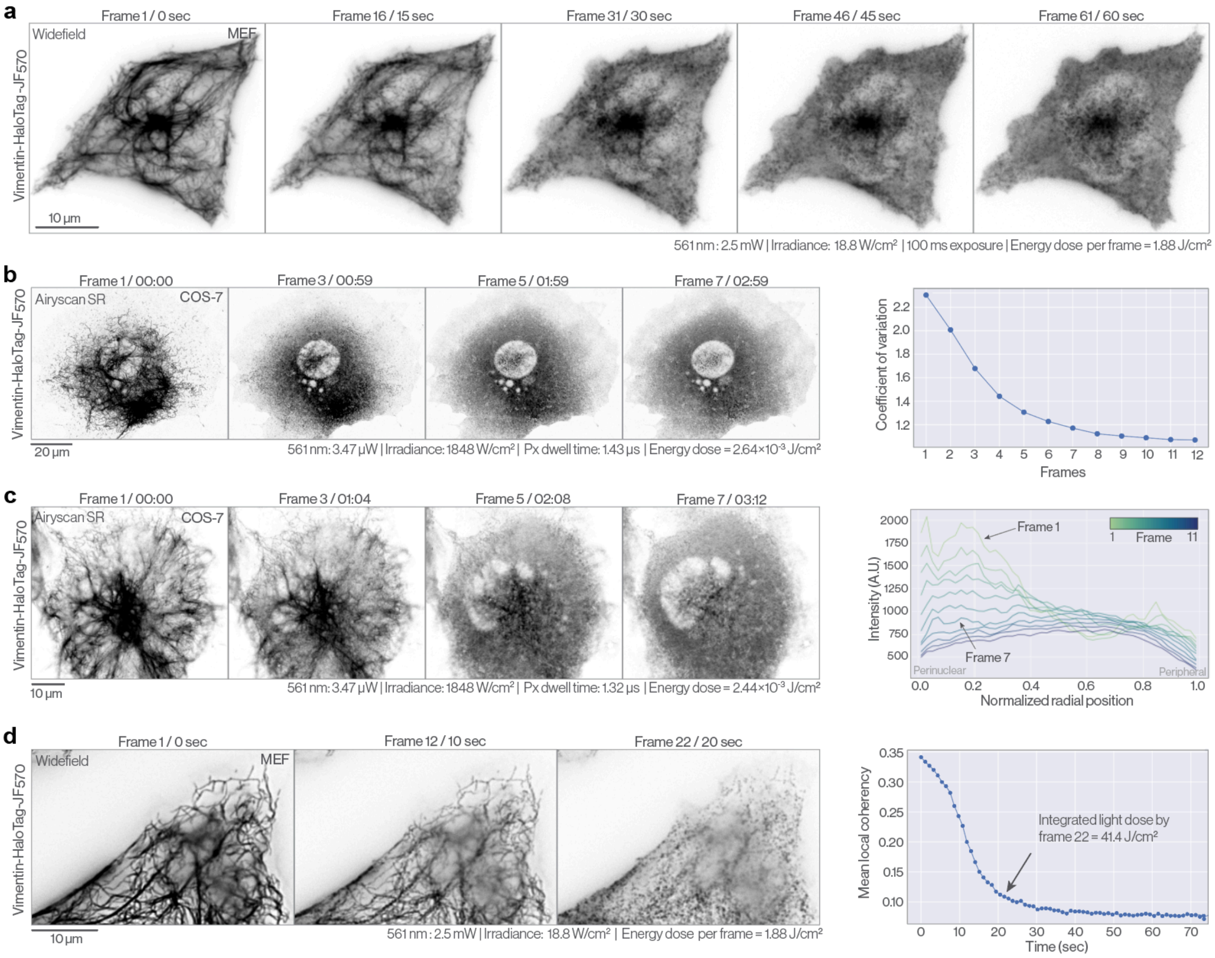


**Supplementary Figure 2: Dynamics of FilaBuster-mediated IF network disintegration**

**a.** Widefield montage of VIF disassembly in a MEF labeled with vimentin-HaloTag-JF$_{570}$. **b.** Airyscan timelapse of VIF fragmentation in a COS-7 cell expressing vimentin-HaloTag-JF$_{570}$. VIF disassembly results in a more homogeneous fluorescence signal throughout the image reflected in the decreasing coefficient of variation over time. **c.** Additional timelapse showing VIF disassembly in a vimentin-HaloTag-JF$_{570}$ labeled COS-7 cell. The mean normalized radial distribution of vimentin intensity between the nuclear envelope (0) and the plasma membrane (1) is plotted over time. Loss of filament integrity is associated with a more uniform distribution of VIF fragments throughout the cytoplasm. **d.** Widefield timelapse of VIF fragmentation in a vimentin-HaloTag-JF$_{570}$ labeled MEF. Light-mediated VIF fragmentation results in the disappearance of dense filament bundles and a steep drop in local orientational coherency within the image. 561 nm laser power, irradiance, pixel dwell time, and energy dose per pixel are reported for confocal experiments a and b. Laser power, irradiance, exposure time, energy dose per frame, and integrated energy dose to VIF fragmentation are reported for c. See methods for additional details.

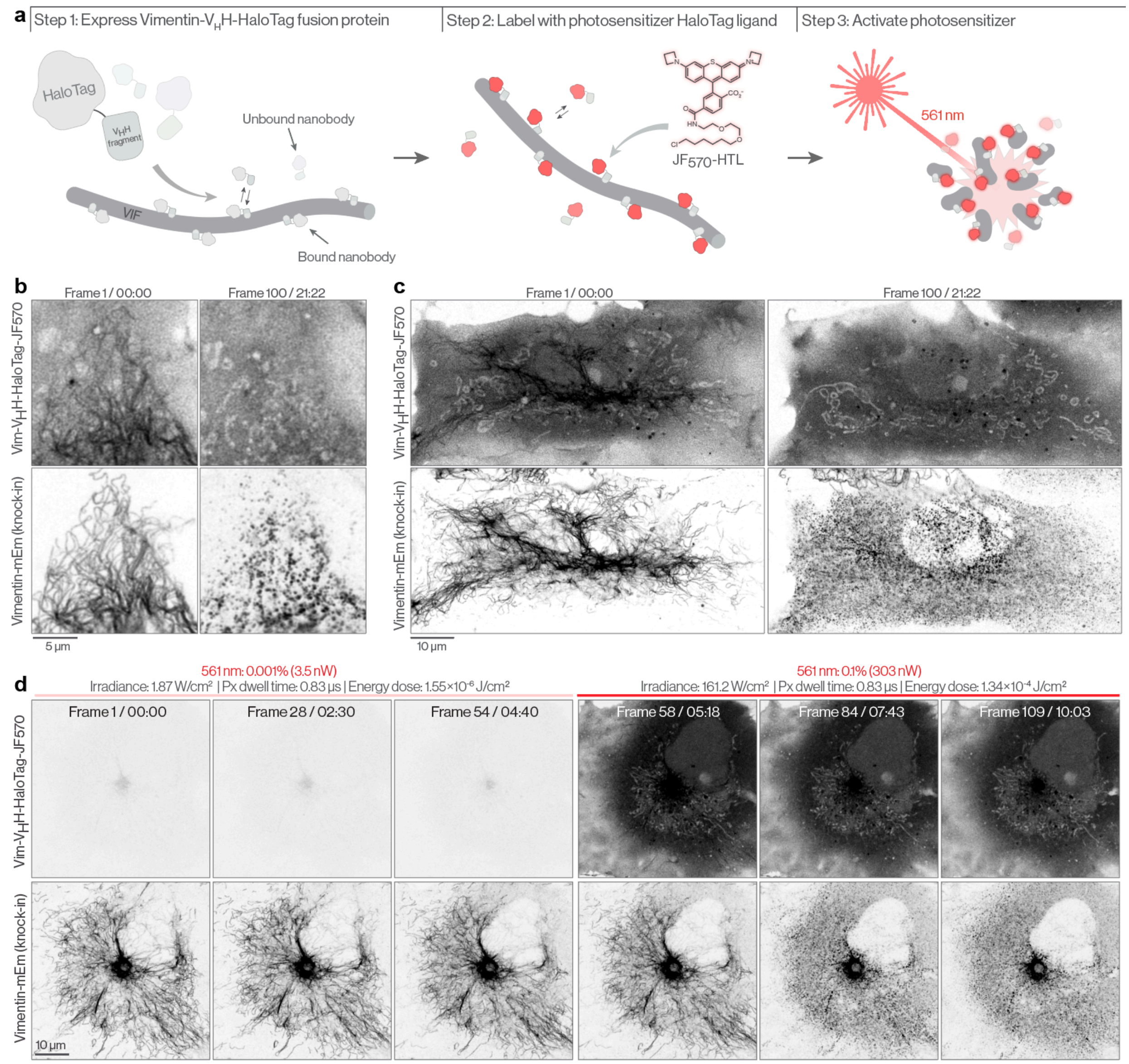


**Supplementary Figure 3: VIF disassembly by activation of a nanobody-targeted photosensitizer**

**a.** Cartoon schematic of the FilaBuster strategy using a Vimentin-VHH/nanobody-HaloTag targeting strategy. **b-c.** Airyscan images of vimentin-$V_H$H-HaloTag-$JF_{570}$ (top) and endogenously tagged vimentin (mEmerald, bottom) in COS-7 cells. After 100 frames / illumination cycles, the vimentin-associated nanobody signal disappears and the endogenous vimentin appears highly fragmented. **d.** Airyscan montage of vimentin-$V_H$H-HaloTag-$JF_{570}$ (top) and vimentin-mEmerald (knock-in, bottom) in a COS-7 cell. Five minutes into the movie (frame 55), the 561 nm laser power is raised two orders of magnitude, activating the photosensitizer and driving vimentin IF disassembly. Irradiance, pixel dwell time, and

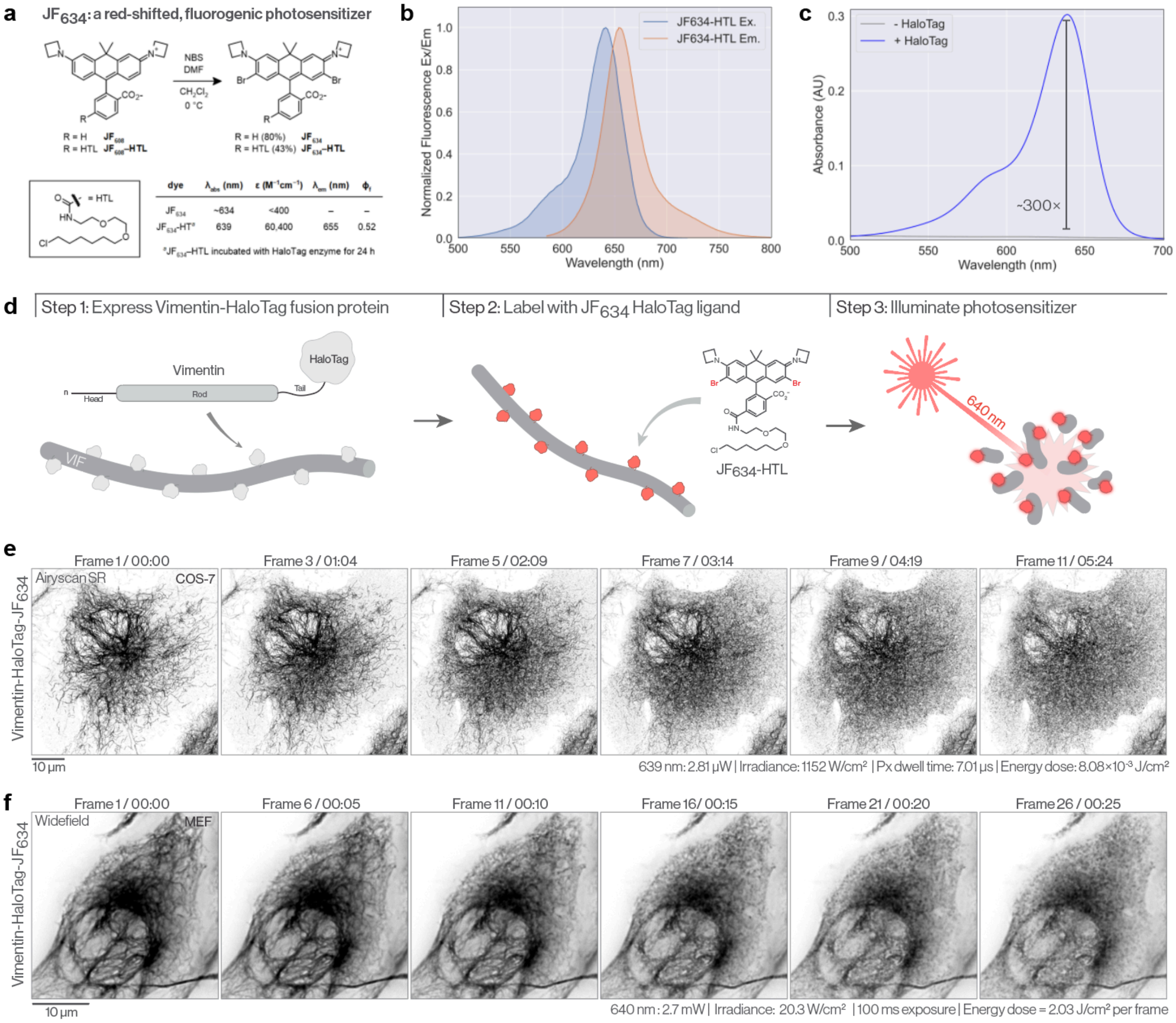


**Supplementary Figure 4: A novel far-red emitting photosensitizer HaloTag ligand**

**a.** Characterization of $JF_{634}$ and $JF_{634}$-HTL. **b.** Excitation/emission spectra for $JF_{634}$-HTL. **c.** $JF_{634}$-HTL is fluorogenic, displaying ~300× greater absorbance upon HaloTag binding. **d.** Cartoon schematic indicating adapted FilaBuster strategy with red-shifted HaloTag ligand and longer wavelength excitation light. **e.** Airyscan montage of a COS-7 cell expressing vimentin-HaloTag-$JF_{634}$ illuminated with 639 nm light (power: 2.8 μW; energy dose: 8.1 $\times 10^{-3}$ J/cm²). **f.** Widefield epifluorescence timelapse of a stable vimentin-HaloTag MEF labeled with $JF_{634}$-HTL.

a

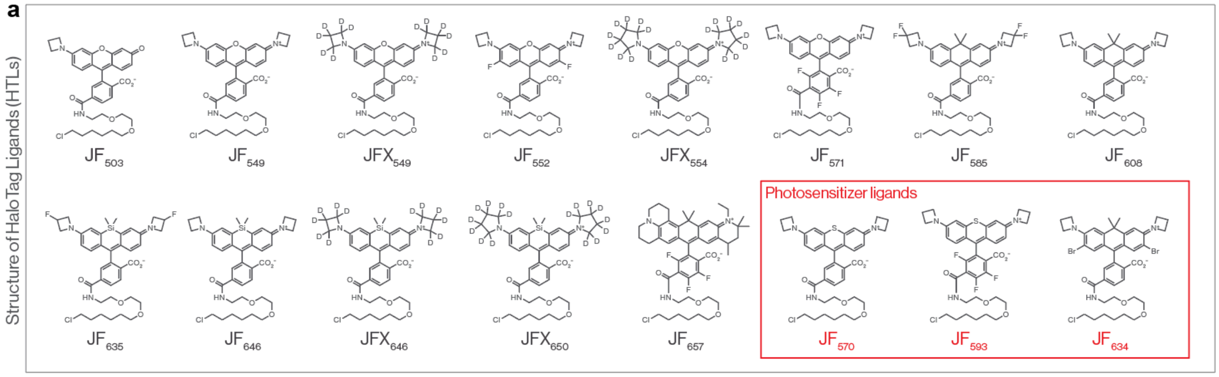


b

| HaloTag Ligand | Laser / power | Irradiance | Energy per frame | Avg. time to VIF fragmentation | Avg. total energy to VIF fragmentation | Sample (cells) |
|---|---|---|---|---|---|---|
| $JF_{503}$ | 488 nm / 2.9 mW | 21.8 W/cm² | 2.18 J/cm² | 95 ± 71 sec | 207.1 J/cm² | 17 |
| $JF_{570}$ | 561 nm / 2.5 mW | 18.8 W/cm² | 1.88 J/cm² | 50 ± 35 sec | 94.0 J/cm² | 34 |
| $JF_{593}$ | 561 nm / 2.5 mW | 18.8 W/cm² | 1.88 J/cm² | 51 ± 30 sec | 95.9 J/cm² | 39 |
| $JFX_{554}$ | 561 nm / 2.5 mW | 18.8 W/cm² | 1.88 J/cm² | 73 ± 27 sec | 137.2 J/cm² | 28 |
| $JF_{549}$ | 561 nm / 2.5 mW | 18.8 W/cm² | 1.88 J/cm² | 86 ± 58 sec | 161.7 J/cm² | 39 |
| $JFX_{549}$ | 561 nm / 2.5 mW | 18.8 W/cm² | 1.88 J/cm² | 90 ± 57 sec | 169.2 J/cm² | 15 |
| $JF_{552}$ | 561 nm / 2.5 mW | 18.8 W/cm² | 1.88 J/cm² | 115 ± 66 sec | 216.2 J/cm² | 45 |
| $JF_{571}$ | 561 nm / 2.5 mW | 18.8 W/cm² | 1.88 J/cm² | 244 ± 169 sec | 458.7 J/cm² | 22 |
| $JF_{608}$ | 561 nm / 2.5 mW | 18.8 W/cm² | 1.88 J/cm² | 312 ± 164 sec | 586.6 J/cm² | 22 |
| $JF_{585}$ | 561 nm / 2.5 mW | 18.8 W/cm² | 1.88 J/cm² | 341 ± 126 sec | 641.1 J/cm² | 63 |
| $JF_{634}$ | 640 nm / 2.7 mW | 20.3 W/cm² | 2.03 J/cm² | 17 ± 19 sec | 34.5 J/cm² | 50 |
| $JF_{646}$ | 640 nm / 2.7 mW | 20.3 W/cm² | 2.03 J/cm² | 61 ± 27 sec | 123.8 J/cm² | 12 |
| $JFX_{646}$ | 640 nm / 2.7 mW | 20.3 W/cm² | 2.03 J/cm² | 81 ± 31 sec | 164.4 J/cm² | 34 |
| $JF_{635}$ | 640 nm / 2.7 mW | 20.3 W/cm² | 2.03 J/cm² | 166 ± 71 sec | 337.0 J/cm² | 48 |
| $JF_{608}$ | 640 nm / 2.7 mW | 20.3 W/cm² | 2.03 J/cm² | 192 ± 59 sec | 389.8 J/cm² | 23 |
| $JF_{657}$ | 640 nm / 2.7 mW | 20.3 W/cm² | 2.03 J/cm² | 202 ± 177 sec | 410.1 J/cm² | 53 |
| $JFX_{650}$ | 640 nm / 2.7 mW | 20.3 W/cm² | 2.03 J/cm² | 271 ± 177 sec | 550.1 J/cm² | 20 |

c

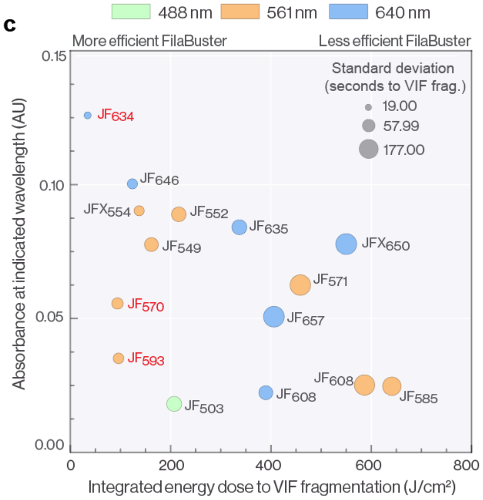


**Supplementary Figure 5: Benchmarking JF-HTL FilaBuster efficiency in vimentin-HaloTag MEFs**
**a.** Structures of HaloTag ligands tested, including conventional rhodamine-based dyes (black) and photosensitizer dyes (red). **b.** Summary table of illumination parameters and performance metrics for each ligand in vimentin-HaloTag MEFs using widefield epifluorescence microscopy. All experiments were performed using a 63×/1.4 NA oil objective, 100 ms exposure time, and 1 frame per second acquisition. Shown are laser wavelength used and power measured at the sample plane, irradiance, energy dose per frame, average time to VIF fragmentation (equivalent to number of illumination cycles), and average total energy dose to fragmentation. **c.** Integrated energy required for VIF disassembly plotted against ligand absorbance at the laser line used for excitation. Bubble color indicates excitation wavelength; bubble size reflects the variability in fragmentation latency. This dataset serves as a practical reference for implementing FilaBuster across different ligands and imaging platforms. Note: $JF_{608}$ was screened with both 561 nm and 640 nm laser lines. Photosensitizer dyes are indicated in red.

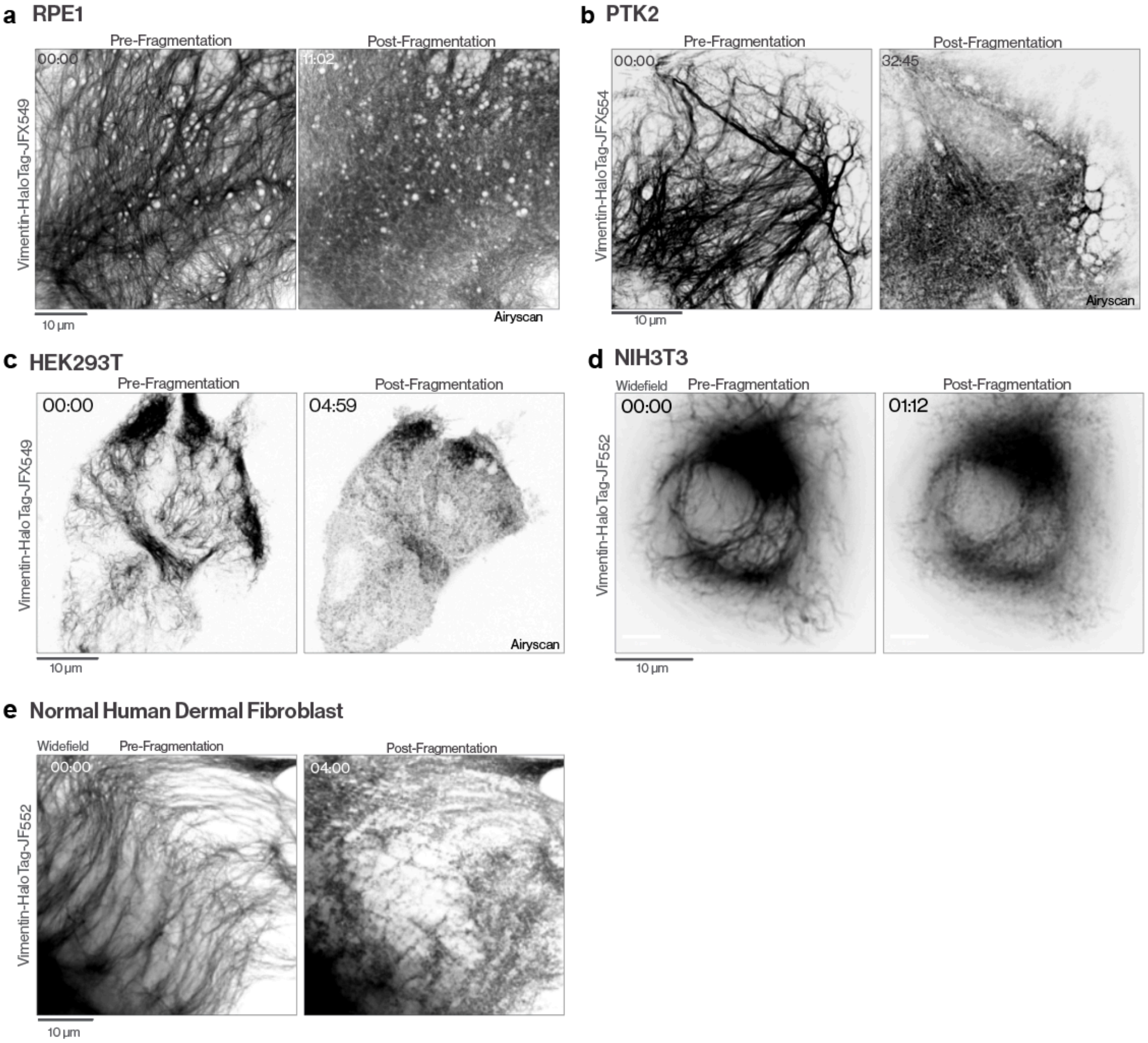


**Supplementary Figure 6: FilaBuster-mediated VIF disassembly is reproducible across cell types**
**a.** RPE1 expressing vimentin-HaloTag-JFX$_{549}$ before (left) and after (right) 561 nm irradiation. **b.** PTK2 cell expressing vimentin-HaloTag-JFX$_{554}$ before (left) and after (right) 561 nm irradiation. **c.**HEK293T cell expressing vimentin-HaloTag-JFX$_{549}$ before (left) and after (right) 561 nm irradiation. **f.** Endogenously tagged vimentin-HaloTag in a U2-OS cell before (left) and after (right) 642 nm irradiation.**d**. NIH3T3 cell expressing vimentin-HaloTag-JF$_{552}$ before (left) and after (right) 561 nm irradiation. **e.** Normal human epidermal fibroblast expressing vimentin-HaloTag-JF$_{552}$ before (left) and after (right) 561 nm irradiation.

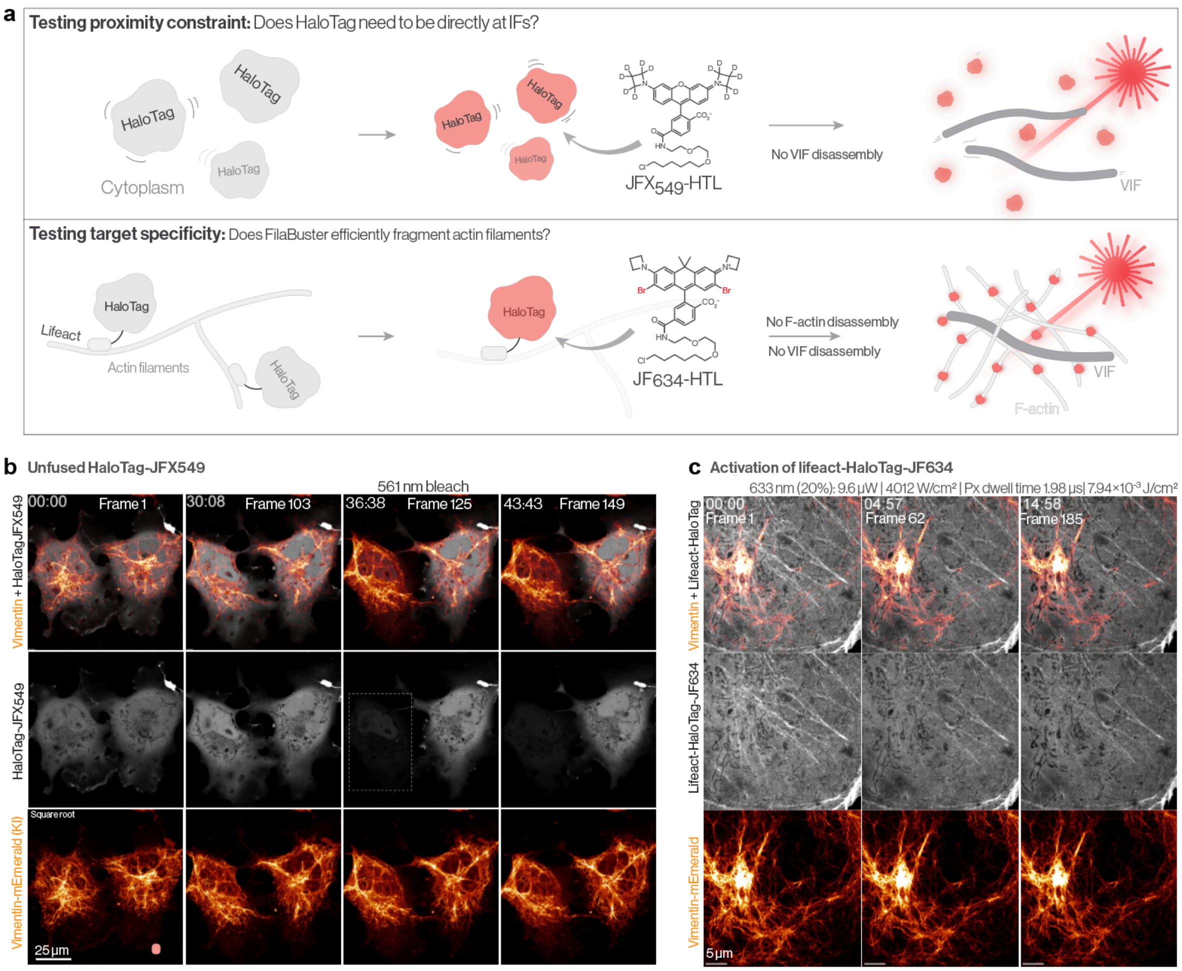


**d** Comparison with light doses used for CALI

| Study | Fusion protein | Chromophore | Effect | Threshold illumination energy for effect |
|---|---|---|---|---|
| Keppler and Ellenberg, 2009 | SNAP-tag-α-Tubulin | BGAF SNAP-tag ligand | Metaphase arrest | 55,000 J/cm² |
| Rajfur et al., 2002 | EGFP- α-actinin | EGFP | Stress fiber detachment from focal adhesions | 105,200 J/cm² |
| Vitriol et al., 2008 | EGFP-Capping Protein | EGFP | Formation of dorsal protrusions | 15,000 J/cm² |
| Tour et al., 2003 | Cx43-TC | ReAsH/FlAsH | Gap junction inactivation | 425-450 J/cm² |
| Takemoto et al., 2011 | HaloTag7-PKCγ | Eosin HaloTag ligand | PKCγ translocation to the plasma membrane | 792 J/cm² |
| Binns et al., 2020 | EGFP-HaloTag | JF$_{570}$-HaloTag ligand | Decreased EGFP fluorescence | 100 J/cm² |
| Current work | Vimentin-HaloTag | JFX$_{554}$-HaloTag ligand | IF fragmentation | 137.2 J/cm² |
| Current work | Vimentin-HaloTag | JF$_{634}$-HaloTag ligand | IF fragmentation | 34.5 J/cm² |

**Supplementary Figure 7: Defining FilaBuster operational boundaries and failure modes**
**a.** Cartoon schematic indicating the two boundary conditions that we investigated: expression of a non-directed HaloTag protein and expression of an F-actin-directed HaloTag. **b.** Airyscan timelapse of endogenously tagged vimentin (vimentin-mEmerald, orange) and unfused HaloTag-JFX$_{549}$ (gray) in a COS-7 cell. At frame 125, the left cell is extensively photobleached with 561 nm light, but we observe no effect on VIF stability seven minutes later. **c.** Testing target specificity. Airyscan montage of lifeact-HaloTag-JF$_{634}$ and vimentin-mEmerald in a COS-7 cell. Prolonged irradiation of the f-actin-targeted photosensitizer does not result in global actin filament fragmentation or loss of VIF integrity. **d.** Table comparing FilaBuster (current work) with relevant publications using various configurations of chromophore assisted light inactivation (CALI).

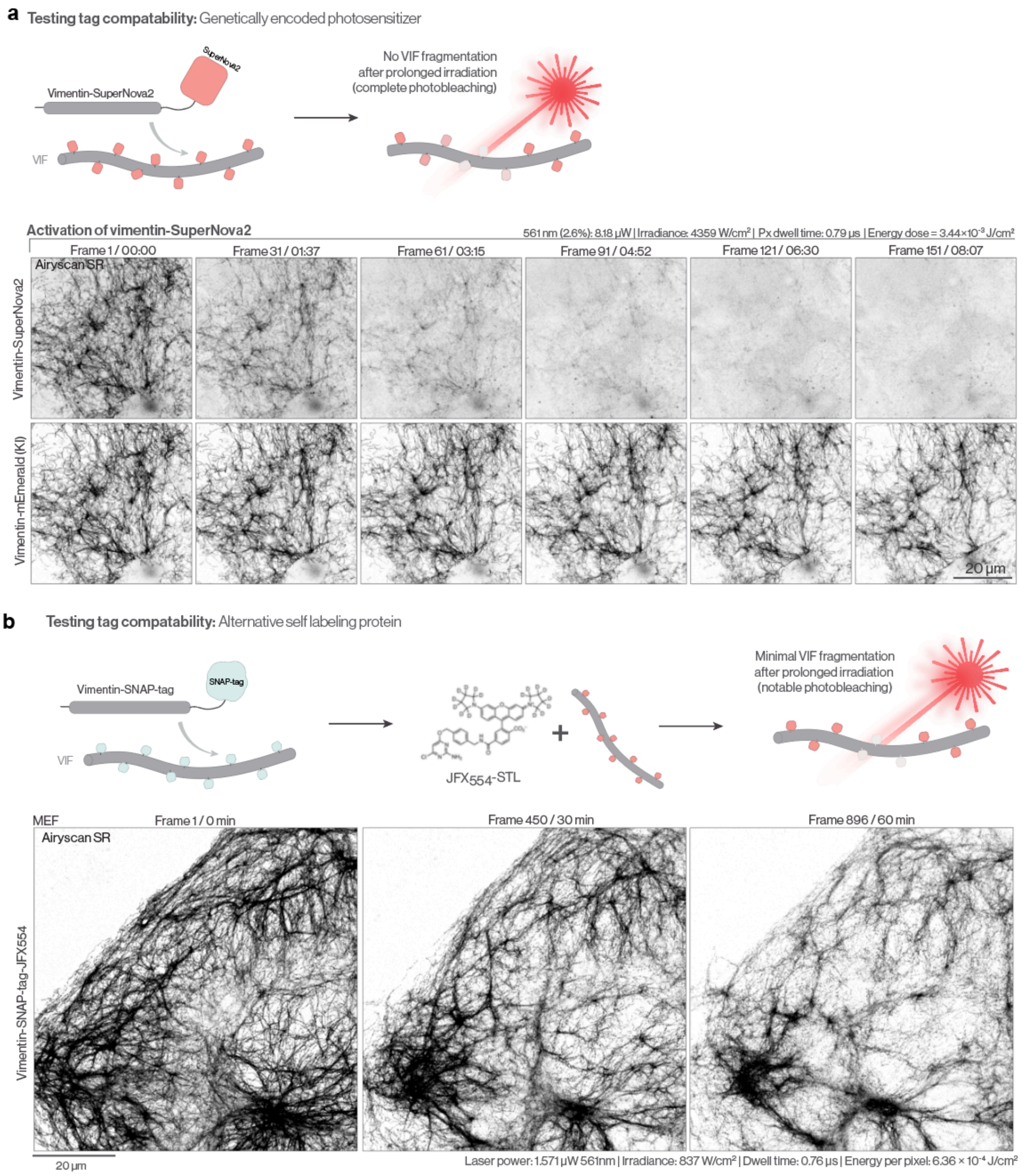


**Supplementary Figure 8: SuperNova2 and SNAP-tag fusions do not support efficient light-mediated VIF disassembly**

**a.** Testing the compatibility of a genetically encoded photosensitizer fluorescent protein. Airyscan montage of vimentin-SuperNova2 (top) and endogenously tagged vimentin (mEmerald, bottom) in a COS-7. After ~60 cycles of 561 nm illumination, the SuperNova2 tag fully photobleaches but we observe no VIF destabilization. **b.** Testing the compatibility of SNAP-tag, an alternative self labeling protein. Airyscan timelapse of a MEF stably expressing vimentin-SnapTag-JFX$_{554}$ displayed negligible VIF fragmentation over 896 cycles of 561 nm illumination.

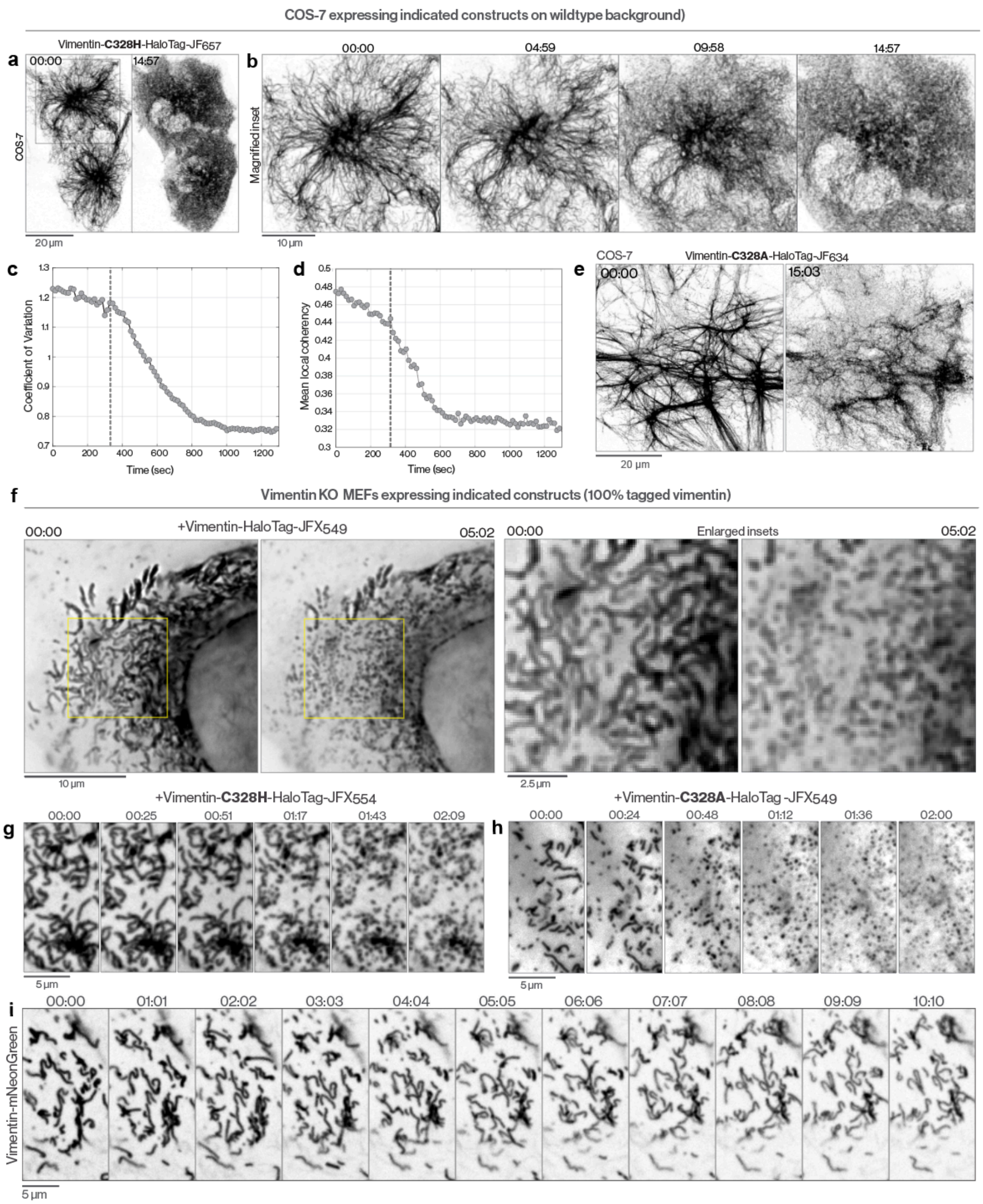


**Supplementary Figure 9: Vimentin C328 is dispensable for FilaBuster-mediated VIF fragmentation**
**a**.Airyscan micrographs of stable COS-7 cells expressing vimentin-C328H-HaloTag-$JF_{657}$ before (left) and after (right) VIF disintegration. **b**. Enlarged inset from A showing montage of VIF disassembly. **(c-d)**. Coefficient of variation (**c**) and average local coherency (**d**) indicating time-course of VIF disintegration over the full movie shown in b. **e**. Airyscan micrographs of a COS-7 cell stably expressing vimentin-C328A-HaloTag-$JF_{634}$ before (left) and after (right) VIF fragmentation. (**f-h**) Vimentin knockout MEFs rescued with either (**f**) Vimentin-HaloTag-$JFX_{549}$, (**g**) vimentin-C328H-HaloTag-$JFX_{554}$, or (**h**) vimentin-C328A-HaloTag-$JFX_{549}$. Expression of 100% tagged vimentin produces truncated ~3 µm filaments which undergo clear disintegration in both control and C328 mutants, confirming that FilaBuster-mediated VIF fragmentation does not require cysteine. (**i**) Vimentin knockout MEF rescued with the conventional vimentin-mNeonGreen fusion. Expression of 100% tagged vimentin-mNeonGreen results in comparable truncated filaments, which do not undergo fragmentation.

**a**

| Sequence | Modifications | # PSMs | Positions in Master Proteins | Theoretical MH+ [Da] | RT [min] | Oxidation Site | Abundances in A | Abundances in B | Abundances in C |
|---|---|---|---|---|---|---|---|---|---|
| [R].LGDLYEEEMR.[E] | 1xOxidation [M9] | 7 | 146-155 | 1270.56 | 40.08 | M154 | 161.3 | 101.4 | 37.2 |
| [R].LQDEIQNMK.[E] | 1xOxidation [M8] | 2 | 365-373 | 1134.55 | 20.63 | M372 | 167.6 | 61.8 | 70.7 |
| [K].LQEEMLQR.[E] | 1xOxidation [M5] | 2 | 189-196 | 1062.52 | 22.91 | M193 | 185.8 | 72.7 | 41.5 |
| [R].EMEENFAVEAANYQDTIGR.[L] | 1xOxidation [M2] | 9 | 346-364 | 2202.96 | 78.41 | M347 | 150.4 | 66.1 | 83.5 |
| [K].MALDIEIATYR.[K] | 1xOxidation [M1] | 1 | 391-401 | 1311.66 | 79.19 | M391 | 171.6 | 36.1 | 92.2 |

The median abundance of peptides with oxidized methionine is 164 under condition A, compared to 69 under condition B and 77 under condition C.

**b**

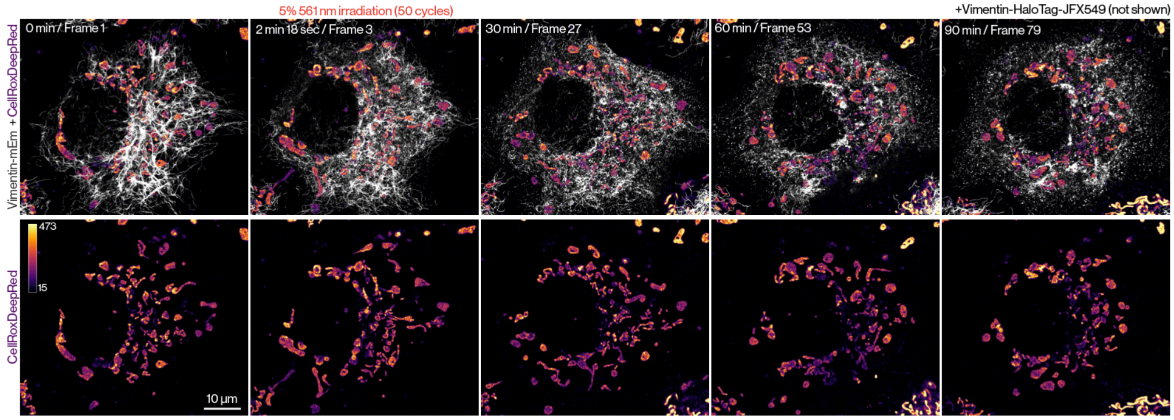


**Supplementary Figure 10: FilaBuster drives vimentin methionine oxidation but minimal global oxidative stress**

**a**. Table indicating differential vimentin methionine oxidation in vimentin-HaloTag knock-in U2-OS cells under three conditions. In condition A, cells were labeled with Biotin-$JF_{646}$-HaloTag ligand and irradiated with 639 nm light to drive VIF fragmentation. In condition B, cells were dye labeled but were not irradiated. In condition C, cells were neither labeled nor irradiated. **b.** Airyscan montage of a vimentin knock-in (mEmerald, white) COS-7 expressing Vimentin-HaloTag-$JFX_{549}$(not shown). The cell was labeled with CellRox DeepRed to monitor changes to global cellular ROS levels. This probe primarily localizes to mitochondria and we observed no detectable change in its intensity in the 1.5 hours after VIF disassembly.

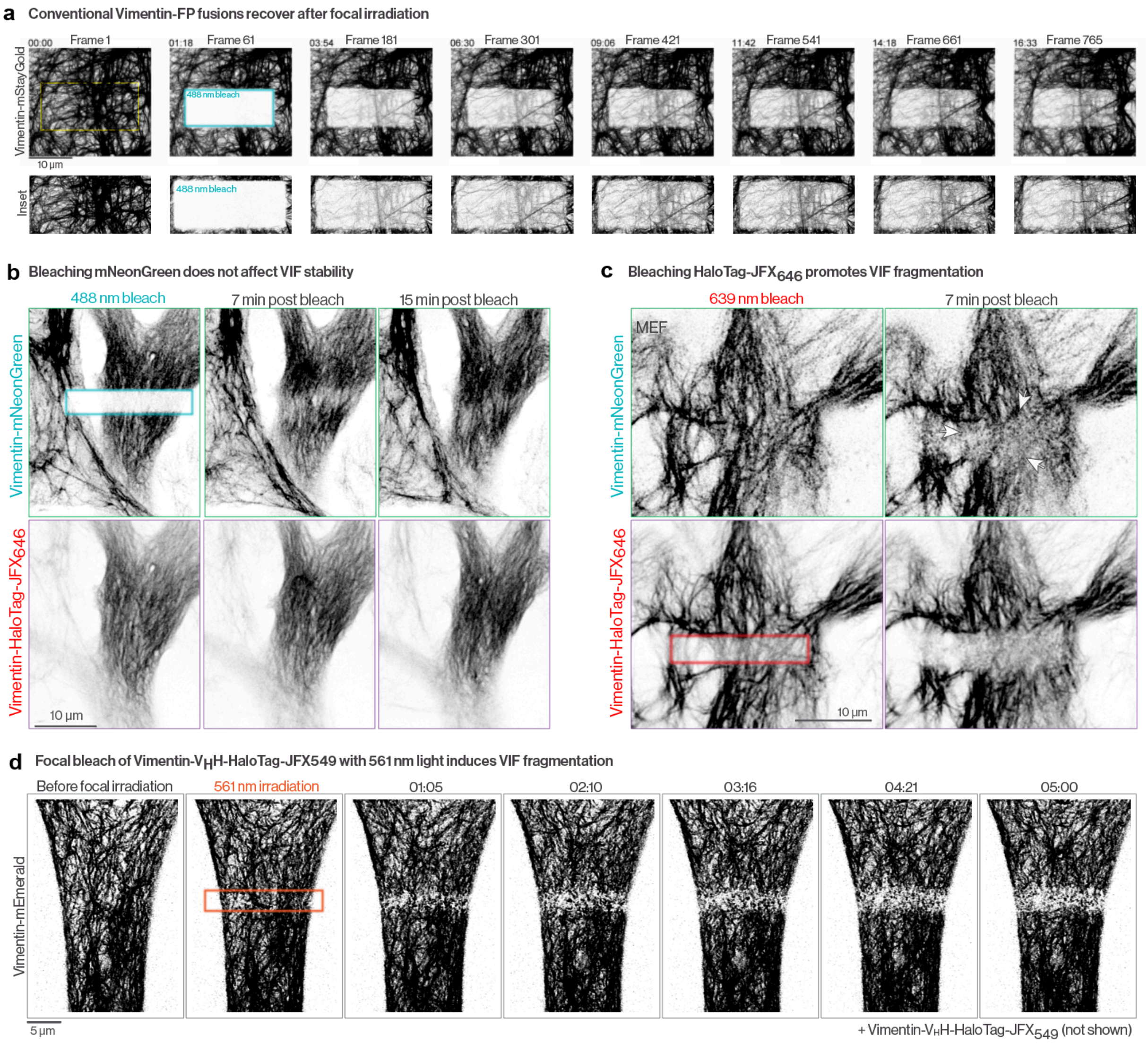


**Supplementary Figure 11: Focal irradiation of conventional vimentin-FP fusions does not induce VIF fragmentation.**

**a.** Airyscan timelapse of vimentin-mStayGold. At frame 61, StayGold is photobleached at the indicated region. Over the subsequent 15 minutes, there is extensive recovery due to trafficking of new filaments into the bleach site and no evidence of VIF fragmentation. **b-c.** MEFs co-expressing vimentin-HaloTag-JFX$_{646}$ with the conventional reporter vimentin-mNeonGreen. **b.** After photobleaching mNeonGreen with 488 nm light, we observe no evidence of VIF fragmentation. **c.** After photobleaching JFX$_{646}$ with 639 nm light, we observe robust VIF fragmentation visible in the mNeonGreen channel as well (arrows). **d.** Airyscan montage of a U2-OS co-expressing the conventional reporter vimentin-mEmerald along with vimentin-V$_H$H-HaloTag-JFX$_{549}$ (not shown). 561 nm irradiation of the JFX$_{549}$-labeled nanobody at the indicated region (red box), results in localized vimentin IF disassembly visible with the conventional mEmerald reporter.

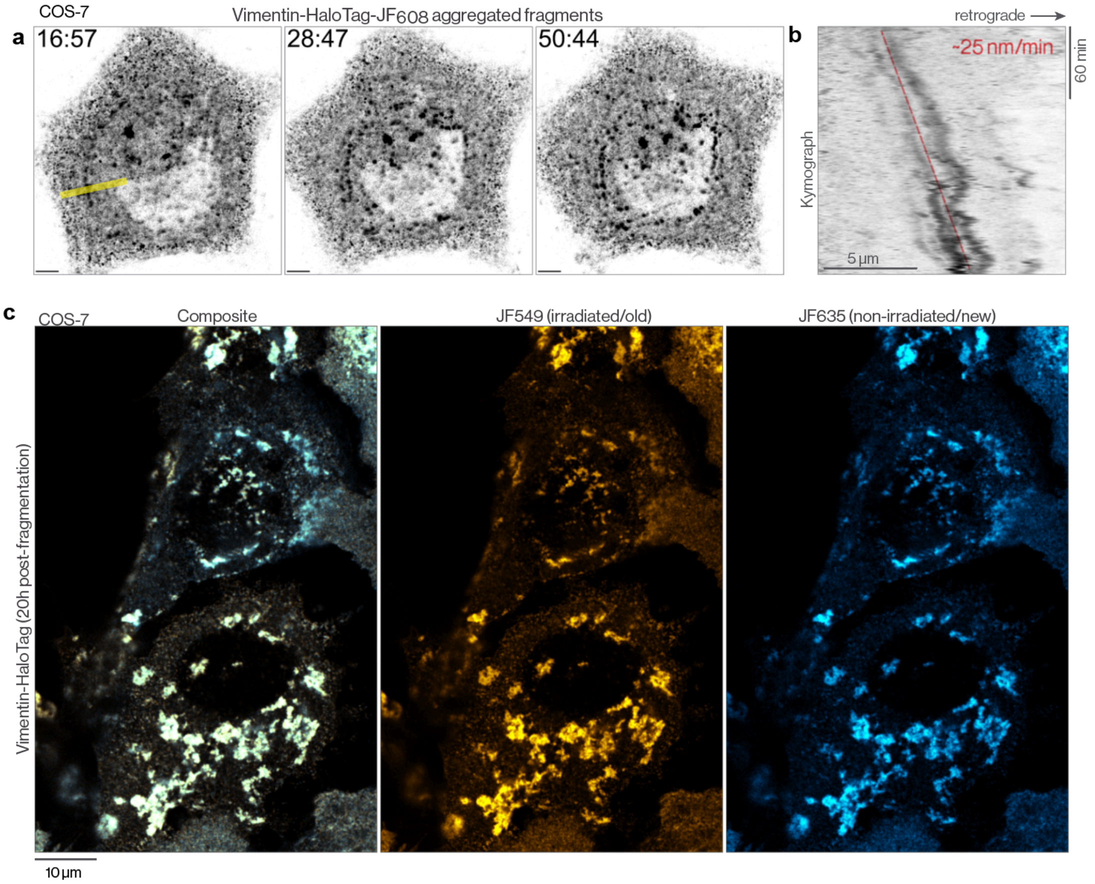


**Supplementary Figure 12: VIF fragments coalesce and inhibit new filament assembly**

**a.** Airyscan montage of vimentin-HaloTag-$JF_{608}$ in a COS-7 at the indicated times post-VIF disassembly (hh:mm). Scale bar is 5 µm. Yellow line indicates line scan used for b. **b.** Kymograph displaying retrograde motion of aggregated VIF fragments over 3+ hours with average retrograde speed of ~25 nm/min. **c.** Pulse chase labeling of vimentin-HaloTag 20 hours after 561 nm light-mediated VIF fragmentation. COS-7 cells were initially labeled with $JF_{549}$-HTL (yellow), extensively washed, and irradiated to drive VIF disintegration. Twenty hours post-irradiation, cells were labeled with $JF_{635}$-HTL (chase dye, blue) for 1 hour to label any newly translated vimentin-HaloTag after VIF fragmentation.

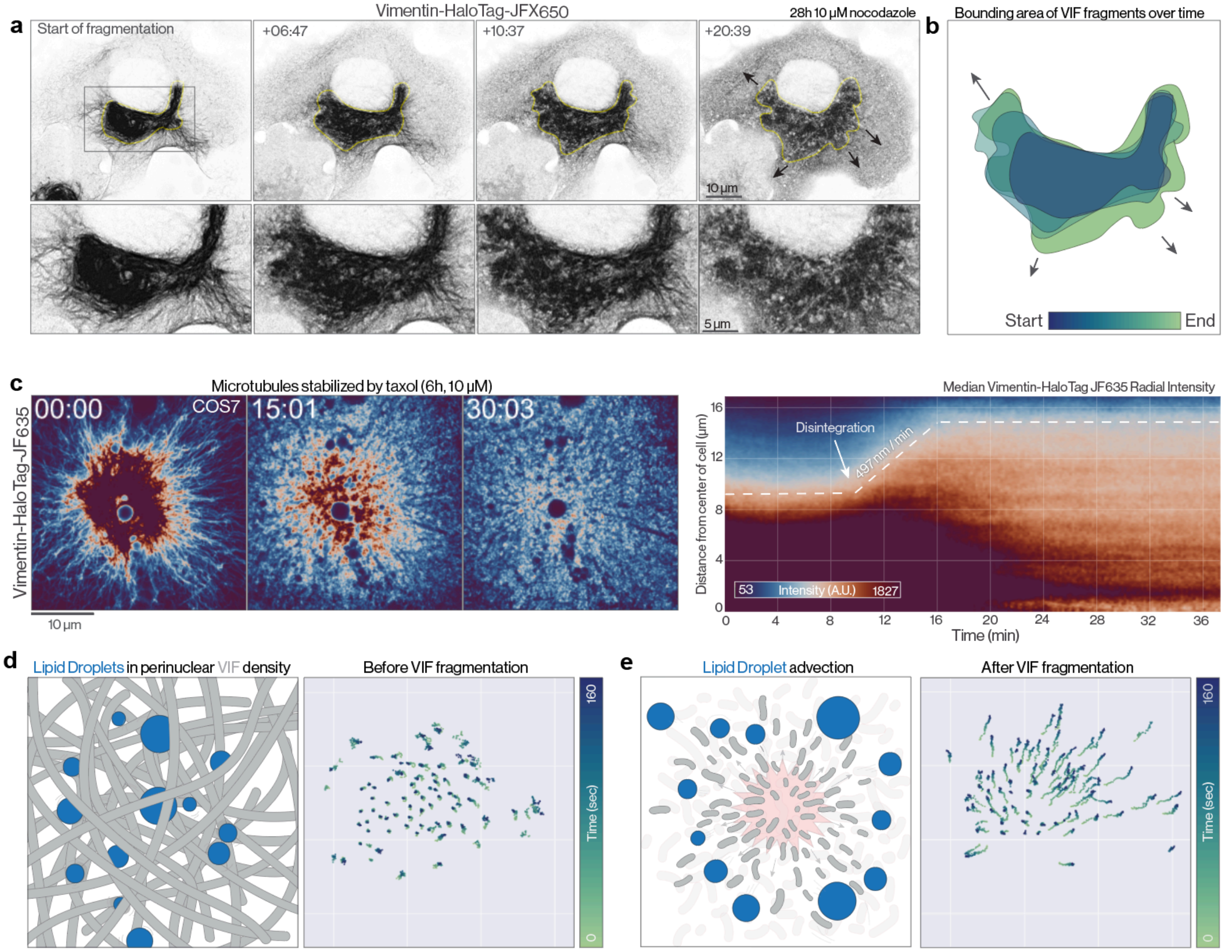


**Supplementary Figure 13: Radial expansion of VIF fragments and perinuclear organelles upon light-induced VIF disassembly**

**a.** Rapid light-mediated disassembly of a collapsed perinuclear VIF network in a nocodazole-treated COS-7 cell. VIF fragments expand outward over 15-20 minutes. **b.** Color-coded bounding area of perinuclear VIF density after 639 nm irradiation indicating radial expansion of VIF fragments. **c.** (left) A radially expanding wave of VIF fragments upon 633 nm irradiation of vimentin-HaloTag-$JF_{635}$ in a taxol-treated COS-7 cell. (right) Kymograph of radial vimentin-HaloTag-$JF_{635}$ intensity over time. Radial intensity distribution at each time point represents the median of 360 radial line scans from the center of the cell, in 1° increments. VIF disintegration at ~10 min, is accompanied by the rapid expansion of VIF fragments traveling at ~497 nm/min. **d-e**. Color coded trajectories of perinuclear, VIF-associated lipid droplets either before (d) or after (e) VIF fragmentation.

**Supplementary video captions:**

**Video 1 | Cryo-tomogram of Vimentin-Halotag from MEF detergent treated cells**
Tomogram movie showing Vimentin filaments (diameter ~11 nm) with associated HaloTags (spherical densities, diameter 3-4 nm) from a MEF ghost cell

**Video 2 | VIF disassembly by activation of vimentin-HaloTag-JF$_{570}$ with 561 nm illumination**
**A.** Airyscan movie of VIF fragmentation in COS-7 expressing vimentin-HaloTag-JF$_{570}$ (see Fig. 1e). **B-C.** Additional examples of VIF fragmentation in COS-7 cells upon 561 nm illumination of vimentin-HaloTag-JF$_{570}$. **D.** Widefield movie of VIF fragmentation in a MEF expressing vimentin-HaloTag-JF$_{570}$ (see Fig. S2d). E. Airyscan timelapse of vimentin-mEmerald (knock-in) in a COS-7 cell co-expressing vimentin-HaloTag-JF$_{570}$ (not shown). VIF fragmentation is induced by 561 nm illumination. Time format: (mm:ss).

**Video 3 | VIF disassembly by activation of a vimentin nanobody-targeted photosensitizer**
**A.** Airyscan movie of vimentin-mEmerald COS-7 expressing vimentin-VHH-HaloTag-JF$_{570}$. At ~5 min, 561 nm laser power is increased to trigger VIF fragmentation (see Fig. S3D). Scale bar = 5 µm. **B-C**. Airyscan movies of VIF fragmentation in COS-7 co-expressing vimentin-mEmerald (knock-in) and vimentin-VHH-HaloTag-JF$_{570}$ (see Fig. S3b-c). Scale bar in B is 5 µm, scale bar in C is 10 µm. Time format: (mm:ss).

**Video 4 | VIF fragmentation upon illumination of vimentin-HaloTag-JF$_{634}$**
**A.** Airyscan timelapse of VIF fragmentation in a COS-7 expressing vimentin-HaloTag-JF$_{634}$ (see Fig. S4e). **B.** Additional example of VIF fragmentation in COS-7 expressing vimentin-HaloTag-JF$_{634}$. Coefficient of variation within the indicated ROI is displayed at each frame. Scale bar B= 5 µm. Time = mm:ss.

**Video 5 | VIF fragmentation using conventional HaloTag ligands**
**A**. Widefield movies of stable vimentin-HaloTag expressing MEFs labeled with different JF-HTLs: i. JF$_{503}$, ii. JF$_{549}$, iii.JFX$_{549}$, iv. JF$_{552}$, v. JFX$_{554}$, vi. JF$_{571}$, vii. JF$_{585}$, viii. JF$_{593}$, ix. JF$_{608}$ (561 nm excitation), x.JF$_{608}$ (640 nm excitation), xi. JF$_{634}$, xii. JF$_{635}$, xiii. JF$_{646}$, xiv. JFX$_{646}$, xv. JFX$_{650}$, xvi-xvii. JF$_{657}$, xviii. JF$_{635}$-biotin, xix. JF$_{669}$. **B.** Airyscan movie of VIF fragmentation in a COS-7 expressing vimentin-HaloTag-JFX$_{650}$ **C.** Lattice SIM movies of VIF fragmentation in COS-7 expressing vimentin-HaloTag labeled with conventional JF-HTLs: i. JFX$_{549}$, ii. JF$_{552}$, iii. JF$_{635}$.

**Video 6 | FilaBuster boundary testing and failure modes**
**A**. Airyscan movies of COS-7 expressing LifeAct-HaloTag-JF$_{634}$ and vimentin-mEmerald **B.** Airyscan timelapse of vimentin-SNAP-tag-JFX$_{554}$ **C.** Airyscan timelapse of vimentin-SNAP-tag-JFX$_{554}$ and vimentin mEmerald (knock-in) in a COS-7. Scale bar = 5 µm. **D.** Airyscan timelapse of vimentin-SuperNova2 and vimentin-mEmerald (knock-in) in a COS-7. Scale bar = 5 µm.

**Video 7 | Localized disassembly of the VIF cytoskeleton**
**A.** Focal disassembly of VIFs in a MEF expressing Vimentin-HaloTag-JFX$_{549}$. Colormap indicates local image orientation, and focal irradiation regions are indicated by circles **B.** Bisection of a perinuclear VIF density by focal 639 nm irradiation of vimentin-HaloTag-JFX$_{650}$ **C.** Focal 639 nm irradiation of multiple ROIs in a COS-7 expressing vimentin-HaloTag-JFX$_{650}$ **D.** Focal 561 nm irradiation of vimentin-HaloTag-JFX$_{549}$ in a vimentin-mEmerald knock-in COS-7 cell. **E.** Focal 561 nm irradiation of a COS-7 expressing vimentin-HaloTag-JF$_{570}$. Whole-cell imaging after targeted irradiation drives global VIF fragmentation due to efficiency of the photosensitizer ligand under standard imaging conditions.

**Video 8 | Localized VIF fragmentation with Vimentin-VHH-HaloTag**
**A.** Severing a VIF bundle with targeted 561 nm irradiation of a vimentin-VHH-HaloTag-JFX$_{549}$. Scale bar = 5 µm. **B.** Vimentin-mEmerald knock-in COS-7 stably expressing vimentin-VHH-HaloTag-JFX$_{549}$. Focal 561 nm irradiation (red circle overlay) induces rapid VIF fragmentation, while focal 488 nm irradiation (green circle overlay) does not **C.** Vimentin-mEmerald knock-in COS-7 + vimentin-VHH-HaloTag-JFX$_{549}$ (not shown). Focal 561 nm irradiation at 01:11 and 06:30 induces fragmentation of targeted VIFs. Close

up of 561 nm bleach region 1 is replayed. **D.** Close-up view of bleach region 1 from (C) showing both vimentin-VHH-HaloTag-JFX$_{549}$ (gray) and vimentin-mEmerald (orange). Scale bar = 5 μm. For all clips, time format: (mm:ss).

**Video 9 | Effects of FilaBuster-mediated VIF disassembly on the actin cytoskeleton**
**A.** Actin filament dynamics in the first 90 minutes after light-induced VIF fragmentation. Airyscan timelapse movies of vimentin-HaloTag-JF$_{635}$ (left) actin filaments (F-Tractin-mCherry, center), and endogenously tagged vimentin (mEmerald, right). At ~6 minutes, the cell is illuminated with 639 nm light, inducing fragmentation of the majority of VIFs. Time format: (hh:mm:ss). **B.** Actin filaments (F-Tractin-mCherry, top) and vimentin (vimentin-mEmerald knock-in, bottom). VIF fragmentation is triggered only in the left cell (arrow) by 639 nm irradiation of vimentin-HaloTag-JF$_{635}$.Time = mm:ss. Scale bar = 10 μm **C.** VIF fragments accumulate on and destabilize stress fibers approximately 90 minutes post FilaBuster. Airyscan time lapse movie of an actin stress fiber (gray, mCherry-F-tractin) and vimentin (orange, vimentin-mEmerald) beginning 1.5 hours after VIF fragmentation by 639 nm irradiation of vimentin-HaloTag-JF$_{635}$ (not shown). VIF fragments accumulate on the surface of stress fibers. Arrows indicate the site of stress fiber destabilization. Time format: (hh:mm:ss) **D.** VIF fragments coalesce into aggregates that are swept towards the nucleus by actin retrograde flow. Airyscan timelapse movie of actin filaments (gray, mStayGold-UtrCH) and vimentin (orange, vimentin-HaloTag-JFX$_{554}$) in the 10 hours after light-mediated VIF fragmentation. VIF fragments coalesce into aggregates that undergo retrograde motion towards the perinuclear region. **E.** VIF fragments (vimentin-HaloTag-JFX$_{549}$) associate with focal adhesion (TagGFP-Paxillin) after FilaBuster

**Video 10 | Effects of FilaBuster-mediated VIF disassembly on the microtubule cytoskeleton**
**A.** Global VIF fragmentation in cells with sparse baseline VIF networks (vimentin-HaloTag-JF$_{608}$) does not destabilize microtubules (EMTB 3xGFP, see Fig. 5c). **B.** Focal VIF fragmentation has minimal effects on microtubules **C.** Global VIF fragmentation in cells with dense baseline vimentin networks results in appreciable microtubule disassembly **D.** VIF fragmentation in a COS-7 pre-treated with taxol to stabilize microtubules **E.** VIF fragmentation in COS-7 cells pre-treated with nocodazole to disassemble microtubules.

**Video 11 | FilaBuster-mediated fragmentation of GFAP, desmin, peripherin, and keratin 18 IFs**
**A.** Timelapse of GFAP IF fragmentation by 561 nm illumination of GFAP-HaloTag-JF$_{552}$ in a COS-7 cell. **B.** Airyscan timelapse of desmin IF fragmentation by 561 nm illumination of desmin-HaloTag-JFX$_{549}$ in a U2-OS cell. **C.** Lattice SIM movie of peripherin IF fragmentation by 561 nm illumination of peripherin-HaloTag-JFX$_{549}$ in a U2-OS cell. **D.** Focal 561 nm irradiation of vimentin-HaloTag-JF$_{552}$ induces local fragmentation of VIFs as well as desmin IFs marked with a co-expressed desmin-EGFP. **E.** Focal 639 nm irradiation of GFAP-HaloTag-JFX$_{650}$ induces local fragmentation of GFAP IFs as well as VIFs marked with a co-expressed vimentin-mApple. **F.** Cytokeratin IF disassembly by 561 nm illumination of keratin 18-HaloTag-JFX$_{552}$ in a COS-7 cell. Close up view shows formation of punctate, granular structures and loss of filament integrity. Scale bar = 5 μm, time = mm:ss for all movies
